# A test-retest reliability and multiverse analysis of denoising pipelines in four common functional MRI task paradigms

**DOI:** 10.64898/2026.11.23.768645

**Authors:** Sahar Uppal, Elizabeth J Davies, Natalie Ertl, Lysia Demetriou, Leon Fonville, Matthew B Wall

## Abstract

Test-retest reliability is a key issue in functional MRI research, yet it is unclear how reliability differs across commonly used task paradigms, and the effect denoising pipelines may have on reliability measures. The present study used four tasks designed to probe a range of cognitive/emotional domains: N-back (working memory), Go/No-Go (response inhibition), Monetary Incentive Delay (MID; reward processing), and OASIS (emotional images) task. Healthy participants (N=25) completed all four tasks in two scanning sessions. Analyses used fMRIPrep for spatial preprocessing. Twelve subsequent denoising pipelines were then implemented, combining head-motion parameters (6 vs. 24), tissue-based regressors, ‘aCompCor’ components and head-motion spike regressors. An additional ‘multiverse’ analysis produced a consensus result across these pipelines. The performance of each pipeline in driving robust statistical results was assessed using residual motion-related variance, mean Z statistics, and the number of suprathreshold (*Z*=3.1, *p*<0.05, cluster-corrected) voxels. Reliability across sessions was quantified using intraclass correlation coefficients (ICC(3,1)), Dice coefficients, and Spearman’s correlations. Denoising pipeline had limited effects on mean Z statistics, but strongly influenced the spatial extent of suprathreshold clusters, particularly on the event-related tasks (Go/No-Go and MID). No single denoising pipeline was shown to be optimal across all tasks, and pipeline effects were present for different contrasts within the same (emotional pictures) task. Inclusion of motion spike regressors did not confer any benefits compared to similar pipelines without spike regressors. The N-back task was the most reliable (ICC: ∼0.45, Dice: ∼0.8, Spearman’s: >0.9) while the Go/No-Go was the least reliable (ICC: ∼0.1, Dice: ∼0.3, Spearman’s: ∼0.6). Group-level reliability measures (Dice, Spearman’s) showed higher values than the individual-level measure (ICC). Tasks producing the strongest activation levels (N-back, emotional images) showed the highest reliability and the least sensitivity to differences in denoising approach. This suggests that task selection, experimental design, and overall experimental power are likely to be more critical determinants of reliability than analytic decisions.

## Introduction

Task-based functional Magnetic Resonance Imaging (fMRI) is a workhorse method in cognitive neuroscience in which stimuli or cognitive tasks are presented, and Blood-Oxygen Level Dependent (BOLD) responses are compared between conditions (e.g. emotional faces vs. geometric shapes, reward vs. loss) to identify neural responses associated with specific task conditions (Friston, Price, et al., 1996). Task-based fMRI is routinely employed in clinical, developmental, and pharmacological studies to examine effects within and between populations (e.g. Ertl et al., 2024; Skumlien et al., 2022; Wall et al., 2025).

Major challenges of task-based fMRI that hinder generalisability relate to reliability, reproducibility, and replicability of results (Bartlett C Frost, 2008). Reliability in this context refers to the consistency of a measurement across repeated examinations. Reproducibility refers to obtaining the same results using the original code/pipeline and dataset, whereas replicability refers to obtaining similar findings in independent datasets using comparable methods. These concepts are distinct but related; a measure which has poor reliability can limit replicability, by increasing measurement variability. Reliability is particularly important when repeated measurements are required, for example in longitudinal studies or within-subject pharmacological studies comparing active drug and placebo treatments. In such cases, poor reliability may obscure genuine effects by increasing the measurement noise floor. FMRI measurements are affected by numerous sources of noise. These include subject-related factors (head movement, fatigue), physiological effects (cardiac and respiratory fluctuations; Kandimalla et al., 2025), and technological sources (scanner instability, image artefacts, and signal drift). Robust preprocessing and accurate modelling of confounding factors are therefore essential for obtaining clean and interpretable data. Eklund et al. (2019) demonstrated the importance of noise modelling in task fMRI by showing that standard parametric inference methods led to inflated false positive rates; however, these were greatly reduced to nominal levels following denoising using ICA-FIX (Salimi-Khorshidi et al., 2014), illustrating how appropriate noise modelling can improve the validity of statistical inference.

The diversity of available denoising strategies is one source of analytical flexibility which can limit the generalisability of fMRI findings. Processing pipelines for fMRI data are complex and require multiple sequential decisions, with each introducing a ‘garden of forking paths’ (Gelman C Loken, 2013) that can substantially alter results. This was demonstrated by Botvinik-Nezer et al. (2020) in the Neuroimaging Analysis Replication and Prediction Study (NARPS), in which 70 independent analysis teams tested nine hypotheses on the same dataset. No two teams used the same analysis pipeline, resulting in substantial variability in the reported findings: for a given hypothesis, the proportion of teams reporting a significant effect ranged from 0-100%. One approach to this problem is ‘fMRIPrep’, a software package that provides a standardised, reproducible and automated approach to many spatial aspects of preprocessing while remaining analysis-agnostic (Esteban et al., 2019). Despite this, considerable flexibility remains downstream, including in temporal and spatial filtering, denoising procedures and statistical modelling.

Multiverse analysis has emerged as one response: rather than committing to a single arbitrary pipeline, many plausible versions of an analysis are run in parallel and a consensus result is derived (Kievit et al., 2022; Mazei et al., 2025; Vuorre C Przybylski, 2024). Because fMRI pipelines have many possible analytical combinations, exhaustive multiverse analysis may be computationally impractical. Recent work has therefore explored more targeted approaches. Dafflon et al. (2022) used a learning-based method to efficiently sample the pipeline decision space rather than testing it exhaustively. Applying a multiverse approach specifically to reliability, Demidenko et al. (2024) tested the effects of smoothing, motion correction, task parameterisation, and contrast selection on reliability of the Monetary Incentive Delay (MID) task (Knutson et al., 2000). They found that overall reliability was low and that all four analytical choices influenced reliability. Importantly, these effects differed between individual- and group-level reliability: analytical choices affected ICC differently from group-level reliability assessed using Spearman similarity. These findings highlight that reliability is not a unitary concept, and that multiple measures may be required to fully evaluate tasks and pipelines.

The choice of denoising method offers substantial analytical flexibility, and there is little consensus in the literature. One common approach is to regress head-motion parameters, either the basic six (three translations, three rotations) or the extended 24-parameter set including derivatives and squares (Friston, Williams, et al., 1996). Another method is to regress signal from anatomical regions assumed to contain little or no BOLD response, typically the white matter (WM) and cerebrospinal fluid (CSF), on the basis that fluctuations in these tissues reflect non-neural effects (Scheel et al., 2022). The ‘aCompCor’ method extends this approach by applying principal component analysis to WM and CSF signals to derive regressors representing their dominant sources of variance (Behzadi et al., 2007; Muschelli et al., 2014). Five components for each tissue have been proposed as a balance between removing noise and retaining task-related signal (Chai et al., 2012). Motion censoring, or ‘spike regression’, involves modelling high-motion volumes as nuisance regressors (Siegel et al., 2013). Data-driven approaches based on Independent Component Analysis (ICA) methods such as ICA-FIX (Salimi-Khorshidi et al., 2014) and ICA-AROMA (Pruim et al., 2015) provide another class of denoising methods, while physiological signals can be regressed out directly with RETROICOR (though these fall outside the scope of the present study; Glover et al., 2000). Across these methods there is an inherent trade-off between removing noise and preserving signal, with potential consequences for reliability. This is further complicated by the loss of temporal degrees of freedom associated with extensive nuisance regression, which can itself bias results (Ciric et al., 2017).

These methods have been extensively evaluated in resting-state fMRI (Ciric et al., 2017; Parkes et al., 2018; Power et al., 2014; Satterthwaite et al., 2019), but comparatively little work has examined their effects in task fMRI, particularly using test-retest reliability. Jones et al. (2022) found that spike regression was generally beneficial, though effects on reliability were modest and often comparable across the pipelines examined. To our knowledge, no study has systematically examined the effect of denoising choices on test-retest reliability across multiple fMRI tasks.

We examined the effects of multiple regression approaches for denoising on signal detection and reliability measures across four commonly used fMRI tasks using data acquired across two sessions. The tasks spanned working memory (spatial N-back), reward processing (MID), response inhibition (Go/No-Go), and emotional processing (positive, negative, and neutral-valenced pictures), comprising two block-designed (N-back, emotion) and two event-related paradigms (MID, Go/No-Go). We compared a set of 12 pipelines comprising different combinations of head motion, WM and CSF signals, aCompCor, and spike regressors, while holding other preprocessing steps constant. We assessed the effectiveness of denoising using FD-DVARS correlation, sensitivity to the experimental effect using Z-values and suprathreshold voxel counts, and reliability using ICC, Dice coefficients, and Spearman’s ρ. We hypothesised that more extensive noise-modelling would improve test-retest reliability, but that additional denoising might yield diminishing returns as complexity increased (Ciric et al., 2017) and could reduce signal-detection measures. Our aim was to identify whether a single denoising pipeline would be optimal across tasks, or whether this is dependent on task and outcome measure.

## Methods

### Participants

Twenty-five right-handed healthy participants (16 females, 9 males) aged 18-50 years (mean age = 25.88 years), participated in this study. All participants were fluent in English. Written informed consent was obtained alongside screening assessments for eligibility. Exclusion criteria were left-handedness, regular medication or psychoactive substance use, history of psychiatric illness, and MRI contraindications. Participants received £10 per scan, with additional monetary compensation based on their MID task performance. Ethical approval was granted by an internal process, and scans were conducted according to Perceptive (London, UK) guidelines and within the Declaration of Helsinki.

### Experimental design

The study consisted of two sessions, both performed at Perceptive Ltd. (Clinical Imaging Centre, Hammersmith Hospital, London), with a separation of 7-14 days. Participants received a training session with demonstration versions of the four fMRI tasks prior to scanning. Tasks were performed in a fixed order: Go/No-Go, OASIS, N-back, MID, with a minimal break between tasks. An operator-guided onscreen reminder of task instructions was presented before each task, and responses were examined to ensure adequate engagement.

### fMRI task paradigms

Tasks were designed based on previous studies (Knutson et al. 2000, Menon et al. 2001, Owen et al. 2005, Kurdi et al. 2017) and presented using PsychoPy (Peirce, 2008). **Figure S1** shows a schematic of each task paradigm.

In the N-back working memory task, participants are required to update their recall of previous trials to respond to each stimulus. Spatial variants of the N-back present stimuli at multiple locations on the screen (Carlson et al., 1998; Owen et al., 2005). A blue square was shown at one of six points on a circle (North, South, Northeast, Northwest, Southeast, Southwest) for 0.6 seconds, with the location changing every 1.5 seconds. In the 0-back block, participants responded whether the stimulus appeared at the fixed target location (North). In the 2-back block, participants responded whether the current stimulus appeared in the same location as the stimulus presented two trials previously. This required constant updating of spatial working memory to maintain the two prior positions. Eight 0-back and eight 2-back blocks were presented, each containing ten trials. Each block lasted 21 seconds, followed by a nine-second rest period. Total task time was eight minutes.

In the MID task, a cue stimulus (an orange or blue square) was presented at the centre of the screen for 0.5 seconds, followed by a 2-4 second delay and then a target stimulus (a white square at the centre of the screen). Participants were instructed to respond to the target stimulus as quickly as possible. The target stimulus duration was initially set at 300 ms and used adaptive timing; if the previous trial was a ‘hit’ (a successful response) then the target duration was decreased by 16.7 ms (one screen refresh). If the previous trial was a ‘miss’ the duration was increased by the same amount. The inter-trial interval was also adjusted to keep trial timings consistent. There were two conditions. In the ‘win anticipation’ condition, the participants earned £1 if the response to the target stimulus occurred while the target was on the screen (a ‘hit’) following the orange cue. In the ‘neutral’ condition, the participant neither won nor lost money based on their response to the target stimulus following the blue cue. The participants received feedback after each trial on their response success and whether they had won money on that trial. Inter-trial intervals ranged from 2-10 seconds (with an approximately Poisson distribution). Twenty-eight neutral trials and 27-win trials were randomly presented throughout the task. Reaction time and money earned were recorded. This task was similar to one used by (Knutson et al., 2000; Skumlien et al., 2022, 2023). Total task duration was 10 minutes and 10 seconds.

The emotional pictures task used the ‘OASIS’ stimulus set by Kurdi et al., (2017); an online and open-source resource that includes 900 colour images with normative ratings on the degree and intensity of emotional valence. Seventy-two images comparable to the International Affective Picture System (IAPS; a standardised database of emotionally evocative images; Lang et al. 2008) were selected. Valence ratings ranged from 1 = ‘very negative’, 2 = ‘somewhat negative’, 3 = ‘neutral’, 4 = ‘somewhat positive’ and 5 = ‘very positive’. These included 24 positive images (mean valence = 5), 24 neutral images (mean valence = 3.5), and 24 negative images (mean valence = 2). Participants were instructed to rate the images’ emotional valence with a five-finger-button response box on the same rating scale. Images were presented for 4 seconds each, with four images in one block (16 s). Images were shown in a pseudo-randomised order with six blocks of each valence type, and six rest blocks. Total task duration was 6 minutes and 24 seconds.

The Go/No-Go task assesses error processing and inhibition (Menon et al., 2001). Participants were instructed to respond to a ‘G’ (go) stimulus as fast as possible by pressing a button and not to respond when the ‘N’ stimulus is presented, a ‘No-Go’ signal. Each stimulus was presented for 0.5 s followed by a variable delay (0.75-2.75 s) and response time was recorded. The absence of response to a go stimulus within 1 s was recorded as an error of omission. Of 210 trials, 168 consisted of Go and 42 No-Go stimuli. Fewer No-Go trials were used to minimise habituation. Total task duration was 8 minutes and 4 seconds.

### MRI data acquisition

All images were acquired on a 3.0 T Magnetom Trio MRI scanner (Siemens Healthcare, Erlangen, Germany) with a 32-channel phased-array head coil. High-resolution anatomical scans were acquired with a magnetisation prepared rapid gradient echo (MPRAGE) T1-weighted sequence (160 slices, TR = 2300ms, TE = 2.98 ms, flip angle = 9°, 1 mm isotropic voxels, bandwidth = 240 Hz/pixel, matrix size = 240 × 256). Functional scans for all tasks were performed with a multiband echo-planar imaging (EPI) sequence for BOLD contrast with 44 axial slices (TR = 1250 ms, TE = 30 ms, flip angle = 65°, 3 mm isotropic voxels, multiband factor = 2, parallel imaging factor = 2, bandwidth = 2232 Hz/pixel). Resting-state scans were also acquired but are not reported here. The functional sequences were based on those tested and validated previously by Demetriou et al. (2018).

### Preprocessing and denoising of fMRI data

Raw fMRI data was converted to Brain Imaging Data Structure (BIDS) format (using the ‘dcm2bids’ software; Boré et al., 2023), for preprocessing in fMRIPrep v24.1.1 (Esteban et al., 2019). See Supplementary Material for the full fMRIPrep output text.

Preprocessing was undertaken within a Singularity container to ensure computational reproducibility by controlling and standardising the software environment. FreeSurfer was used for cortical surface reconstruction and anatomical processing (Fischl et al. 2013); outputs were written to native space and MNI152NLin6Asym space at 2 mm resolution. Anatomical preprocessing included bias field correction, skull stripping, tissue segmentation, surface reconstruction, and spatial normalisation to MNI152NLin6Asym standard space using nonlinear registration. This enabled extraction of physiological noise regressors from WM and CSF, and the other regressors used in the subsequent analysis.

Functional preprocessing consisted of motion correction, registration of functional images to anatomical space, and generation of brain masks and confound time series. Head motion was assessed using framewise displacement (FD; Power et al., 2012). As the aim was to investigate approaches to modelling noise, a liberal exclusion threshold of 1 mm mean FD was set.

The following confound regressors were extracted from the fMRIPrep outputs and used in the subsequent analyses to model non-neural sources of variance in the BOLD signal:

#### Head motion parameters (HMP)

Six head-motion parameters (translations: x, y, z; rotations: pitch, roll, yaw), their temporal derivatives, and squared terms of the original parameters and derivatives produced 24 parameters in total. Head-motion is a well-established confound in fMRI, as it can systematically bias activation estimates and reduce reliability (Power et al., 2014).

#### Tissue-based physiological regressors

Mean signal time series were extracted from WM and CSF masks and used as nuisance regressors. As these regions are assumed not to contain neural signals of interest, their mean signal reflects non-BOLD fluctuations, such as cardiac and respiratory signals, head-motion, and other noise sources (Power et al., 2014). These regressors are commonly used to denoise both task (Ertl et al., 2024; Mills et al., 2023) and resting-state (Ertl, Freeman, et al., 2024; Wall et al., 2019, 2025) data.

#### Anatomical component-based noise correction (aCompCor)

This approach uses WM and CSF signals but performs principal component analysis (PCA) on the time series to derive multiple regressors from each tissue class (Behzadi et al., 2007). It offers a data-driven elaboration of the basic tissue-based mean signal regression by modelling multiple orthogonal components of signal variance (Muschelli et al., 2014). The top five components for WM and CSF were used, totalling 10 parameters.

#### Spike regressors

Volumes exhibiting excessive motion or signal change were modelled using spike regressors to prevent high motion timepoints from biasing parameter estimates. This approach preserves the full timeseries while ‘censoring’ volumes affected by high levels of motion to reduce their impact on contrast estimates (Siegel et al., 2013). The threshold for identification of problematic volumes was FD = 0.5mm (the default in fMRIPrep).

#### Denoising pipelines

From these four sets of variables, 12 sets of denoising regressors were generated; these are described in **Table 1**.

**Table 1.** List and description of the denoising pipelines used in this study.

| **Pipeline** | **Definition** |
|---|---|
| **6HMP** | Basic six head motion parameters. |
| **24HMP** | Six head motion parameters, their temporal derivatives, the squares of the basic six, and the squares of the temporal derivatives. |
| **6HMP-WM-CSF** | Basic six head motion parameters; WM and CSF regressors. |
| **24HMP-WM-CSF** | Twenty-four head motion parameters; WM and CSF regressors. |
| **6HMP-aCompCor** | Six head motion parameters; ten aCompCor regressors (five derived from WM, and five derived from CSF). |
| **24HMP-aCompCor** | Twenty-four head motion parameters; ten aCompCor regressors. |
| **6HMP-spike-reg** | Six head motion parameters; motion spike regressors. |
| **24HMP-spike-reg** | Twenty-four head-motion parameters; motion spike regressors. |
| **6HMP-WM-CSF-spike-reg** | Six head motion parameters; WM and CSF regressors; motion spike regressors. |
| **24HMP-WM-CSF-spike-reg** | Twenty-four head motion parameters; white matter and CSF regressors; motion spike regressors. |
| **6HMP-aCompCor-spike-reg** | Six head motion parameters; ten aCompCor regressors; motion spike regressors. |
| **24HMP-aCompCor-spike-reg** | Twenty-four head motion parameters; ten aCompCor regressors; motion spike regressors. |

*Note*: WM = white matter, CSF = cerebrospinal fluid.

These combinations are non-exhaustive but reflect options commonly used in previous work, while avoiding redundant combinations (e.g. combining WM/CSF with aCompCor regressors).

For each participant, session, and task, the relevant regressors were extracted from the fMRIPrep confound files and combined into 12 denoising regressor sets which were saved as text files for subsequent analysis.

### First-Level analyses

First-level statistical analyses were performed in FSL’s FEAT (v6.0) using a general linear model framework. Task-specific explanatory variables (EVs) were defined using event timing text files. A 100 s high-pass temporal filter and prewhitening were applied to account for temporal autocorrelation, after which pipeline-specific regressors were incorporated into the design matrix. Spatial smoothing was applied using a 6 mm FWHM Gaussian kernel. Registration was disabled in FEAT as the functional data were already normalised. The following task-specific contrasts were specified as appropriate: 2-back > 0-back (N-back), No-Go > Go (Go/No-Go), Win > Neutral (MID), and positive > neutral and negative > neutral (OASIS). Across the four tasks, 12 preprocessing pipelines, and 50 scan sessions, 2400 first-level models were calculated.

### Mid-Level analysis

Mid-level analyses were conducted in FEAT using first-level results from each session. For each task and preprocessing pipeline these were entered into a within-subject paired design (fixed effects) with a session regressor alongside regressors for each subject. This produced subject-specific contrast maps reflecting the mean activation across both sessions. One mid-level analysis was conducted for each of the 12 pipelines and the four tasks. The resulting contrast images from these mid-level models formed the inputs for group-level analyses and selected reliability analyses described below.

### Group-Level analyses

To visualise mean activation across participants, the outputs of the mid-level analyses were entered into a one-sample group-level model in FSL FEAT (v6.0) with inference performed using the FLAME1 model. The resulting statistical maps were thresholded using cluster-based correction (*Z=3.1*, cluster-corrected *p*<0.05).

Session-specific group analyses were also performed to characterise mean task-evoked activation within each session per pipeline. First-level FEAT directories were used in a one-sample group-level model in FSL FEAT separately for each task session, task, and pipeline, using FLAME1. Resulting statistical maps were thresholded using the same cluster-correction (*Z=3.1*, cluster-corrected *p*<0.05). The resulting thresholded maps were used in the appropriate reliability measures below.

### Multiverse analysis

To generate a consensus ‘multiverse’ result across all the pipelines for each task, the unthresholded group-level *Z*-score activation maps for each pipeline were averaged. These averaged *Z*-maps were then thresholded using FSL’s ‘easythresh’ function (*Z=3.1*, cluster-corrected *p*<0.05).

### Pipeline comparisons: residual variance

To quantify the effectiveness of each pipeline at removing variability related to head motion, the relationship between head motion, quantified using framewise displacement (FD), and residual temporal signal variability, quantified using DVARS, was assessed using Spearman’s rho. DVARS was calculated as the root mean square of the change in voxel-wise fMRI signal between consecutive volumes. Pearson’s correlations were first calculated between FD and HRF-convolved task regressors to assess linear collinearity between head motion and the experimental design.

A nuisance-only general linear model was applied to participants minimally preprocessed functional data, and DVARS was calculated from the resulting residual time series. As task regressors were not included in this model, task-related variance remained in the residuals. FD was matched with the later DVARS volume pair before calculating Spearman’s rho separately for each session. For spike-regression pipelines, observations corresponding to detected motion spikes and the immediately subsequent volume were excluded. Session-level correlation coefficients were averaged to obtain one FD-DVARS correlation coefficient per subject, task, and pipeline.

Across subjects, FD-DVARS measures were summarised using the median, interquartile range (IQR), and Fisher Z-transformed mean. For each subject-task combination, the twelve denoising pipelines were ranked according to their FD-DVARS Spearman’s rho, from 1 (lowest rho) to 12 (highest rho), with tied values assigned the average rank. Ranks were subsequently aggregated across subjects within each task and across all subject-task combinations to compare relative pipeline performance.

### Pipeline comparisons: statistical cost

Temporal degrees of freedom (tDOF) loss was quantified to characterise the cost of each nuisance model. Each nuisance regressor was treated as consuming one temporal degree of freedom, and tDOF loss was expressed as a percentage of the total number of volumes acquired in each run. For spike-regression pipelines, the number of nuisance regressors varied according to the number of detected motion spikes. Session-level tDOF loss percentages were averaged across the two sessions to obtain subject-level estimates. Overall pipeline estimates were calculated by pooling subject-level tDOF loss estimates across all tasks and summarising the resulting distribution using the median and IQR.

### Pipeline comparisons: measures of sensitivity

The mean *Z*-statistic and number of suprathreshold voxels were extracted from each thresholded group-level statistical map for each task, contrast, and pipeline using ‘fslstats’ (FSL v6.0). These measures were chosen as robustness and spatial extent of activation clusters are commonly reported in fMRI studies.

### Pipeline comparisons: measures of test-retest reliability

Voxelwise ICC(3,1) was calculated for each task, contrast, and preprocessing pipeline using the pyrelimri library in Python (Demidenko et al., 2024). For each computation, functional images from session 1 and 2 were paired across participants and submitted to voxelwise ICC analysis across the whole brain. Additionally, ICC values were calculated within task-specific image masks defined separately for each task and derived from meta-analytic data using Neurosynth (Yarkoni et al., 2011; https://neurosynth.org/). These were selected by the terms ‘working memory’, ‘inhibitory control’, ‘monetary incentive’ and ‘emotional processing’ for the N-back, Go/No-Go, MID and OASIS tasks respectively (Figure S10).

ICC estimates, 95% confidence intervals, and between-/within-subject variance maps were saved as CSVs and NIfTI images. ICC was calculated as:

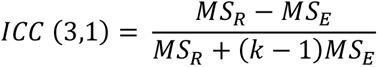

where *MS_R_* is the mean square for rows (between subjects), *MS_E_* is the mean square error (residual) and *k* is the number of measurements (sessions).

Summary statistics including mean, median, SD, and number of voxels included, were also calculated. Ninety-five percent confidence intervals were computed as mean ± 1.96 × (SD / √N), where N is the number of suprathreshold voxels.

Spatial overlap of activation maps between sessions was quantified using Dice coefficients for each task, denoising pipeline, and contrast. Cluster-corrected thresholded *Z*-statistic maps from Session 1 and 2 (*Z=3.1*, cluster corrected *p*<0.05) were binarised and voxelwise intersections between sessions were computed (using fslmaths and fslstats). The Dice coefficient was calculated as:

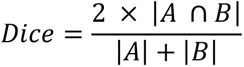

where |A| and |B| denote the number of suprathreshold voxels in Session 1 and 2, and ∣A∩B∣ denotes the number of active voxels common to both sessions. This yields a value between 0 (no spatial overlap) and 1 (perfect overlap), enabling direct comparison of test-retest spatial reliability.

### Spearman correlation

Test-retest reliability of unthresholded activation patterns was assessed using the Spearman rank correlation coefficient, a spatial metric similar to the Dice coefficient. However, unlike Dice coefficients, this approach uses full continuous *Z*-statistic maps and is therefore insensitive to threshold choice. The same meta-analytic masks used in the ICC analysis were used to restrict analyses to functionally meaningful brain regions. Spearman rank correlations were computed using the scipy.stats.spearmanr function from SciPy (Python v3.9) for each task, pipeline and session pair, as:

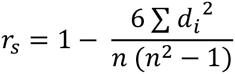

Where *d_i_* denotes the difference in ranks between Session 1 and 2 for voxel *i*, and *n* denotes the number of voxels within the mask. In practice, implementation accounted for tied ranks.

The resulting coefficient ranged from −1 to 1, with higher values indicating greater spatial consistency of activation patterns across sessions.

## Results

### Head motion

No subjects/scans were excluded based on the mean FD threshold. Head motion did not significantly differ between the two scan sessions for the N-back, MID, and OASIS tasks (**Table S1)** but there was a significant difference in the Go/No-Go task (*t*[24] = 2.76, *p* = 0.043; Session 1 mean FD = 0.115 mm; Session 2 mean FD = 0.133 mm). See **Figure S2** for head motion plots and **Table S2** for numbers of spike regressors (FD>0.5mm) for each subject, session, and task. Analyses of the relationship between head motion (mean FD) and activation strength (mean *Z-*statistic) showed no significant relationships (**Figure S3**) in any task or scan session.

### Pipeline effects on residual variance

Collinearity between task condition (regressors) and FD was not demonstrated within any task with Fisher mean of Pearsons r range of −0.093-0.071 (Table showing correlations per task condition are shown in **Table S3)**.

Spearman’s rho of FD-DVARS aggregated across subjects produced the smallest association with 24HMP pipelines across tasks, with inclusion of additional nuisance regressors reducing the association depending on the task (**Figure 1**). The best performing (closest to 0 correlation) was 24HMP-spike-reg in the N-back and MID, 24HMP in Go/No-go, and 24HMP-WM-CSF-spike-reg in the OASIS task. In several pipelines, negative associations were induced between FD and DVARS. A table of the median Spearman’s rho FD-DVARS by task and pipeline can be seen in **Table S4**.

**Figure 1.**
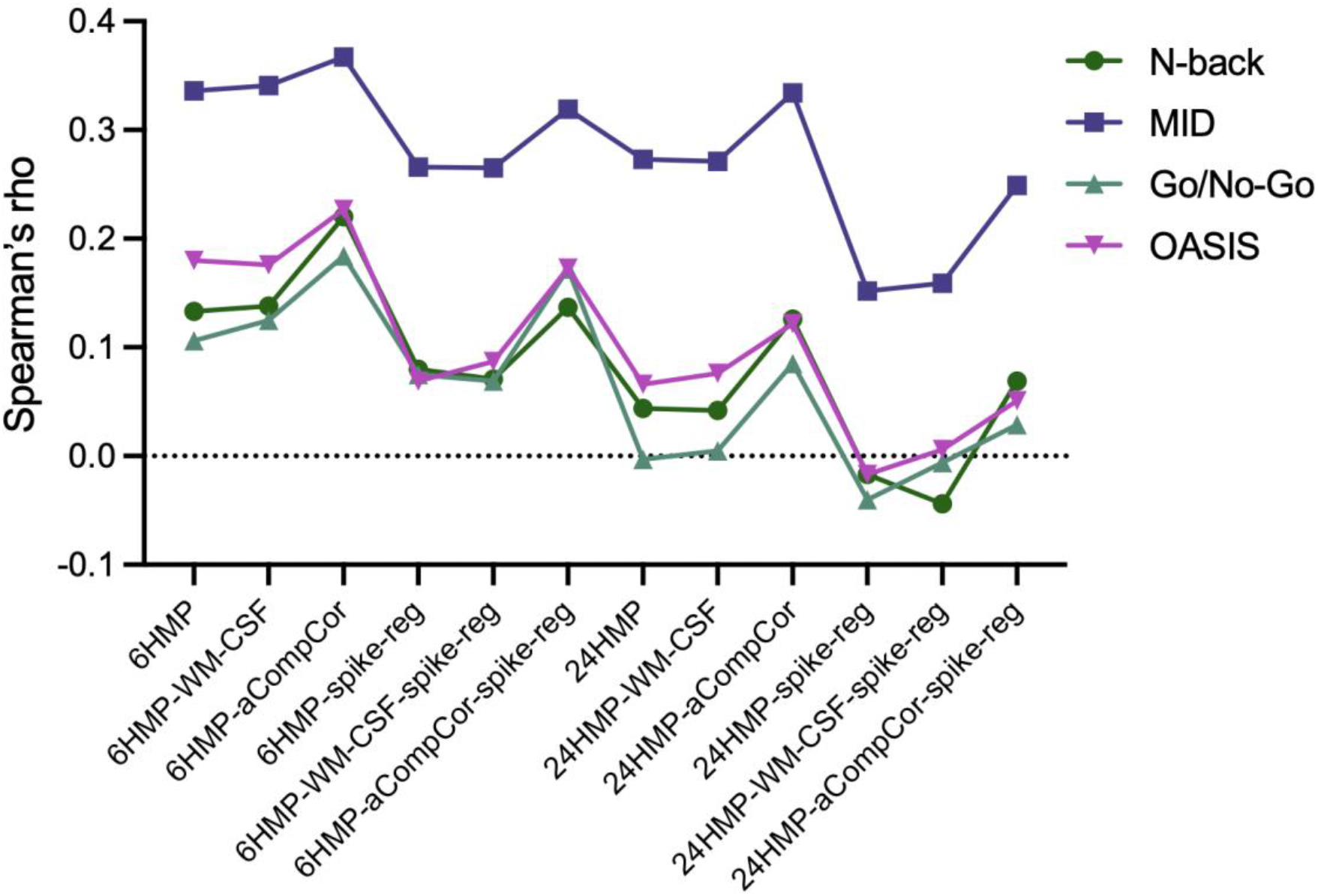
Pipeline effects on the FD-DVARS measure: the correlation between head-movement (FD) and residual variance after nuisance regression (DVARS). Median values across subjects are shown for each task.

When median values were rank transformed, 24HMP-spike-reg and 24HMP-WM-CSF-spike-reg were jointly favoured as the pipelines that best removed motion-related variance, given by the smallest FD-DVARS across participants, sessions and tasks. Summary statistics and ranks per pipeline across tasks are displayed in **Table S5**. Pipelines using aCompCor performed comparatively worse, with 6HMP-aCompCor the most consistently lowest ranked pipeline, appearing in the bottom three ranks 97% of the time.

### Statistical cost

The pattern of tDOF loss reflected the complexity of denoising pipeline with simple pipelines (e.g. 6HMP) reducing the tDOF the least and more complex pipelines (e.g. 24HMP-aCompCor-spike-reg) consuming the most. Results of tDOF loss per pipeline are shown in **Figure S4** and **Table S6**.

### Sensitivity results

Overall activation results (mean of both sessions) are shown in **Figure 2** for each task. These data are derived from the ‘multiverse’ analysis, i.e. the mean of the unthresholded contrast *Z-*statistic images over the 12 pipelines, with the resulting mean subsequently thresholded (*Z*=3.1, cluster-corrected *p*<0.05). Individual activation maps for all 12 pipelines for each task can be found in **Figures S5-SG**. Activation patterns generally accord with previous work using these tasks. For example, the fronto-parietal working memory circuit is evident in the N-back task, the MID task shows strong striatal activation, the Go/No-Go task engaged key inhibitory/cognitive control regions such as the anterior insula and anterior cingulate, and there is strong visual cortex activity in the OASIS task. These results therefore serve to validate the acquisition and analysis procedures used and demonstrate that the tasks produced consistent and expected results.

**Figure 2.**
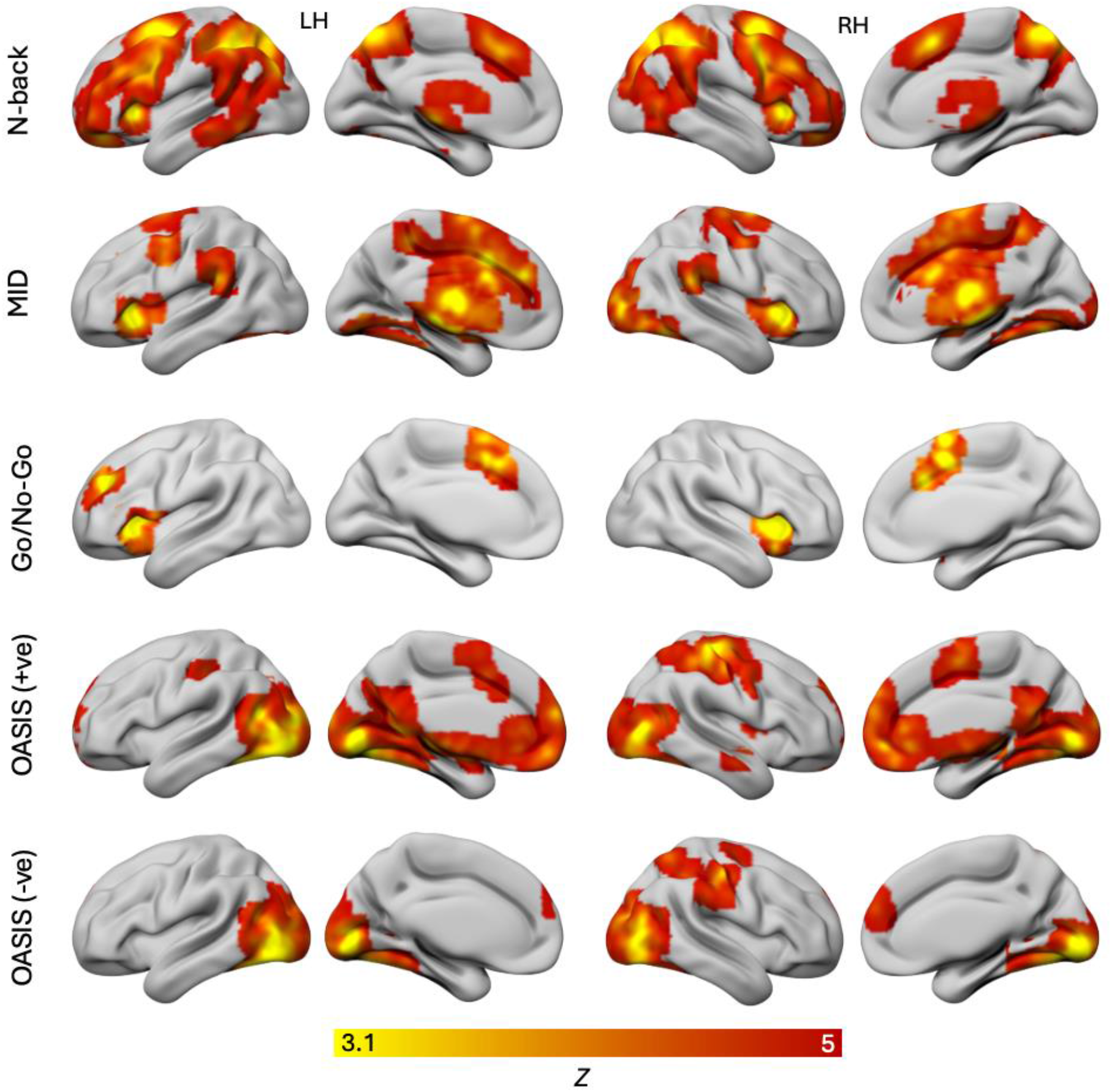
Results from the multiverse analysis (mean of the unthresholded group-level activation maps from the 12 pipelines, *Z*=3.1, *p*_(cluster-corrected)_<0.05). OASIS (+ve) = OASIS positive > neutral and OASIS (-ve) = OASIS negative > neutral.

### Number of suprathreshold voxels

Results from the analysis of suprathreshold voxels in the group-level results for each pipeline are shown below in **Figure 3**. For the N-back task, thresholded activation maps were relatively stable with minimal deviation from the mean across pipelines. In the Go/No-Go task, pipelines using regressors from tissue-based masks (WM/CSF regressors, or aCompCor) performed substantially better than those using purely motion-parameter-derived regressors. The reverse pattern is true for the MID task. The OASIS task showed a more complex pattern with relatively stable performance across all pipelines, but with some variation between contrasts. The negative > neutral contrast showed higher suprathreshold voxel counts in pipelines with WM and CSF regressors, while the positive > neutral contrast showed higher values in pipelines that used aCompCor.

**Figure 3.**
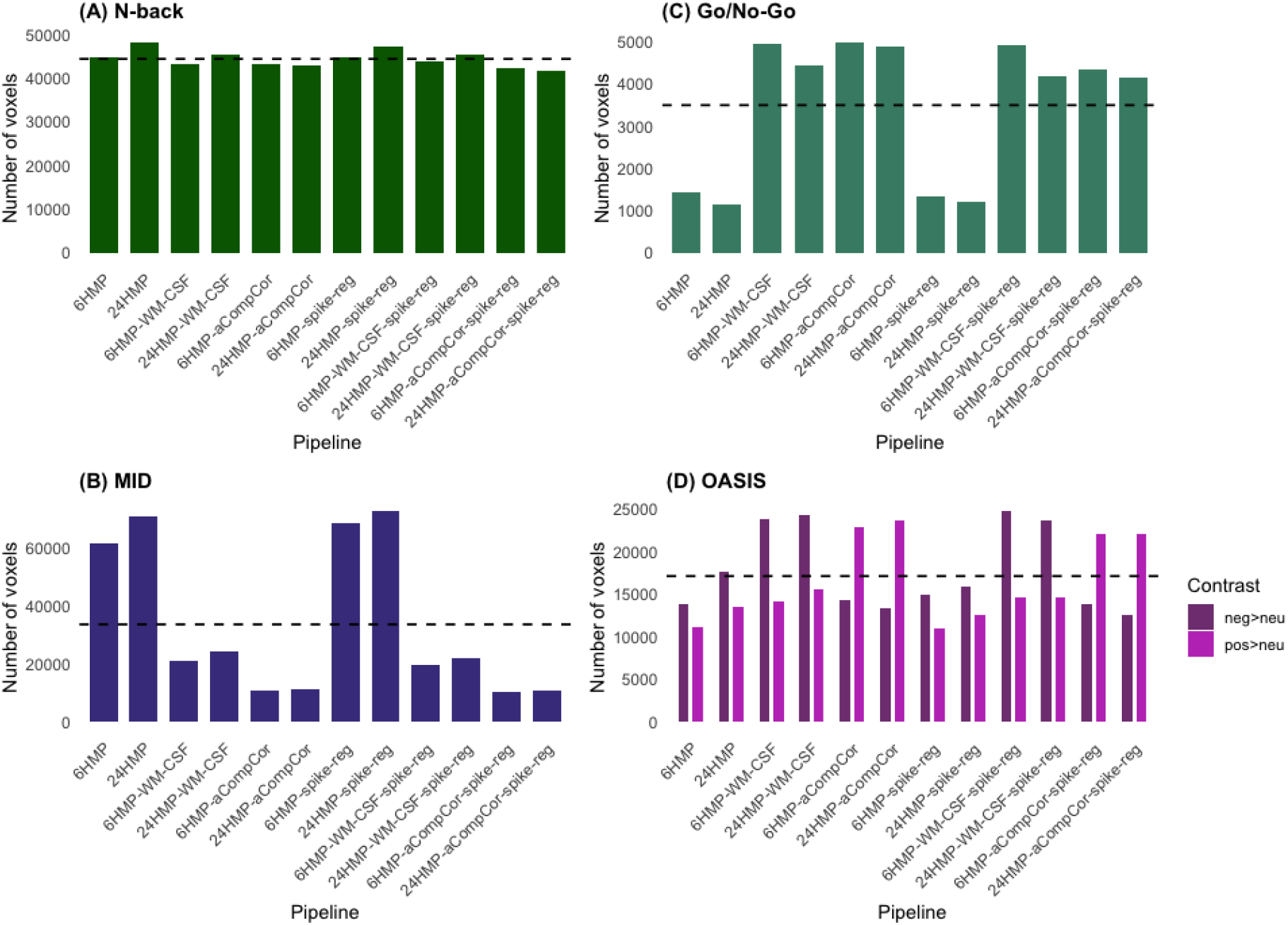
Bar plots displaying the number of suprathreshold voxels across preprocessing pipelines for each task. Results are shown for suprathreshold voxels surviving cluster-level correction *Z=3.1*, *p_(cluster-corrected)_*<0.05 for the (A) Go/No-Go, (B) MID, (C) N-back, and (D) OASIS. Positive > neutral contrast in light purple and negative > neutral contrast in dark purple. The dashed line represents the mean number of suprathreshold voxels across pipelines within each task.

The same data is represented in **Table 2**, alongside the mean *Z-*statistic. Mean *Z-*statistics were relatively stable across pipelines in all tasks, but number of suprathreshold voxels varied by up to a factor of 7 in some tasks (notably the MID task).

**Figure 4.**
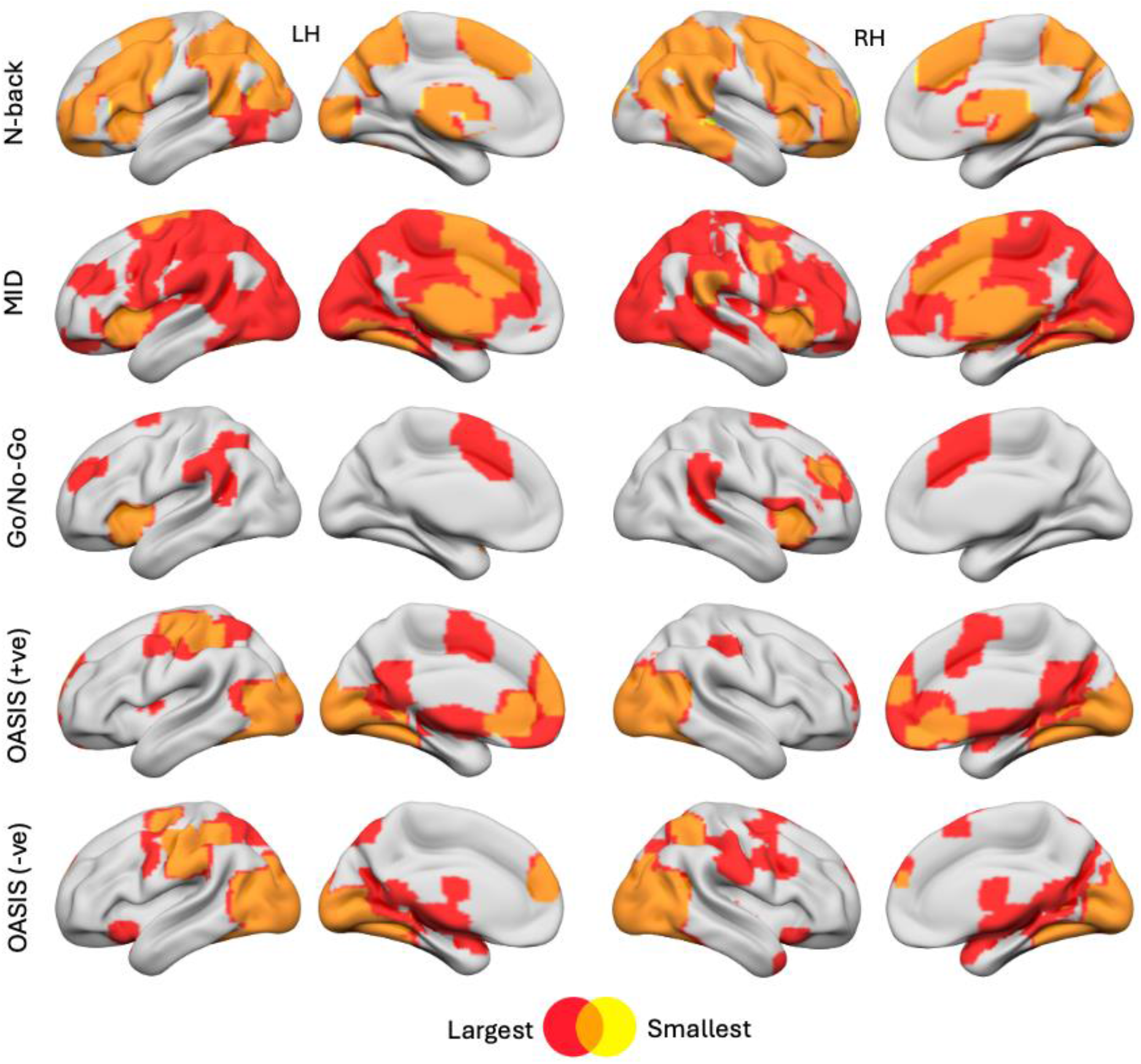
Activation maps projected onto a smoothed cortical surface, showing the pipelines with the smallest (yellow) and largest (red) spatial extent, defined as the number of suprathreshold voxels (*Z*=3.1, *p*_(cluster-corrected)_<0.05), across the 12 pipelines for each task. Orange indicates overlapping regions between the largest and smallest maps. OASIS (+ve) = OASIS positive > neutral and OASIS (-ve) = OASIS negative > neutral. **Table 2** contains details of the pipelines corresponding to the smallest and largest spatial extents for each task.

**Table 2.** Mean *Z*-statistic and number of suprathreshold voxels for each preprocessing pipeline across all pipelines and tasks.

|  | N-back |  | MID |  | Go/No-Go |  | OASIS (pos > neu) |  | OASIS (neg > neu) |  |
|---|---|---|---|---|---|---|---|---|---|---|
|  | Mean *Z* | Voxels | Mean *Z* | Voxels | Mean *Z* | Voxels | Mean *Z* | Voxels | Mean *Z* | Voxels |
| **6HMP** | 4.27 (0.79) | 44626 | 3.78 (0.51) | 61661 | 3.65 (0.42) | 1441 | 4.03 (0.65) | 11094 | 4.01 (0.67) | 13756 |
| **24HMP** | 4.29 (0.81) | **48179** | 3.79 (0.50) | 72122 | 3.68 (0.45) | 1144 | 3.99 (0.67) | 13435 | 4.01 (0.69) | 17623 |
| **6HMP-WM-CSF** | 4.27 (0.78) | 43089 | 3.74 (0.49) | 20876 | 3.69 (0.48) | **4943** | **4.06** (0.70) | 14116 | 4.00 (0.70) | 23768 |
| **24HMP-WM-CSF** | 4.28 (0.79) | 45552 | 3.72 (0.47) | 24472 | 3.68 (0.47) | 4414 | 4.02 (0.69) | 15582 | 3.97 (0.69) | 24273 |
| **6HMP-aCompCor** | 4.26 (0.78) | 43095 | 3.77 (0.52) | 10893 | **3.72** (0.47) | 4973 | 4.05 (0.70) | 22865 | 4.04 (0.69) | 14264 |
| **24HMP-aCompCor** | 4.25 (0.78) | 43903 | 3.74 (0.51) | 11053 | 3.71 (0.47) | 4878 | **4.06** (0.71) | **23608** | 4.01 (0.68) | 13294 |
| **6HMP-spike-reg** | 4.28 (0.8) | 44911 | 3.79 (0.51) | 68497 | 3.63 (0.39) | 1334 | 3.99 (0.65) | 11013 | **4.05** (0.70) | 14972 |
| **24HMP-spike-reg** | **4.30** (0.81) | 47292 | **3.80** (0.50) | **72615** | 3.67 (0.43) | 1214 | 4.01 (0.68) | 12502 | 3.99 (0.68) | 15922 |
| **6HMP-WM-CSF-spike-reg** | 4.29 (0.8) | 43850 | 3.72 (0.47) | 19649 | 3.66 (0.44) | 4898 | 4.02 (0.68) | 14657 | 4.01 (0.72) | **24667** |
| **24HMP-WM-CSF-spike-reg** | 4.29 (0.8) | 45301 | 3.72 (0.46) | 21758 | 3.66 (0.45) | 4172 | 4.02 (0.86) | 14536 | 3.96 (0.68) | 23699 |
| **6HMP-aCompCor-spike-reg** | 4.26 (0.78) | 42439 | 3.75 (0.51) | 10391 | 3.68 (0.43) | 4321 | 4.04 (0.70) | 22023 | 4.03 (0.68) | 13875 |
| **24HMP-aCompCor-spike-reg** | 4.25 (0.78) | 41654 | 3.73 (0.50) | 10647 | 3.70 (0.46) | 4143 | 4.02 (0.68) | 22104 | 3.99 (0.66) | 12582 |

*Note*: Voxels represent those surviving cluster-level correction (*Z=3.1*, *p*<sub>(cluster-corrected)</sub><0.05). Bold values indicate the highest mean Z-statistic and voxel count within each task.

### Dice coefficient analyses

Dice coefficients were used to quantify the spatial overlap of suprathreshold activation maps between sessions, thereby assessing reliability of activation topography independent of effect magnitude. Results are shown in **Table 3**. The N-back task showed approximately 80% overlap of group-level spatial activation maps across sessions. The two event-related tasks were less consistent with Dice coefficients ranging from 0.184 to 0.396 (Go/No-Go) and 0.320 to 0.496 (MID), while the OASIS task had moderate overlap (neg>neu, 0.570-0.702/pos>neu, 0.519-0.714).

**Table 3.** Dice coefficients between Session 1 and 2 for each task and preprocessing pipeline.

|  | N-back | MID | Go/No-Go | OASIS (neg>neu) | OASIS (pos>neu) |
|---|---|---|---|---|---|
| **6HMP** | 0.806 | 0.460 | 0.356 | **0.702** | **0.714** |
| **24 HMP** | 0.808 | **0.496** | 0.354 | 0.641 | 0.676 |
| **6HMP-WM-CSF** | 0.807 | 0.404 | 0.261 | 0.635 | 0.577 |
| **24HMP-WM-CSF** | 0.803 | 0.431 | 0.276 | 0.701 | 0.686 |
| **6HMP-aCompCor** | 0.802 | 0.386 | 0.272 | 0.615 | 0.519 |
| **24HMP-aCompCor** | 0.815 | 0.331 | 0.378 | 0.643 | 0.690 |
| **6HMP-spike-reg** | **0.818** | 0.354 | 0.372 | 0.583 | 0.649 |
| **24HMP-spike-reg** | 0.804 | 0.466 | 0.303 | 0.619 | 0.650 |
| **6HMP-WM-CSF-spike-reg** | 0.813 | 0.320 | **0.396** | 0.633 | 0.676 |
| **24HMP-WM-CSF-spike-reg** | 0.815 | 0.335 | 0.378 | 0.570 | 0.657 |
| **6HMP-aCompCor-spike-reg** | 0.806 | 0.381 | 0.184 | 0.646 | 0.554 |
| **24HMP-aCompCor-spike-reg** | 0.801 | 0.374 | 0.264 | 0.632 | 0.536 |

*Note*: Bold values indicate the best performing pipeline for each task.

### Intraclass correlation analyses

To assess spatial reliability of task-evoked activation, ICC(3,1) was calculated voxelwise across the whole brain for each task and pipeline. The resulting ICC maps from the 6HMP pipeline are shown in **Figure 5** as an illustrative example. Values from within task-relevant ROI masks (derived from independent meta-analytic data from https://neurosynth.org/) were then extracted to assess test-retest reliability. These masks are shown in **Figure S10**.

**Figure 5.**
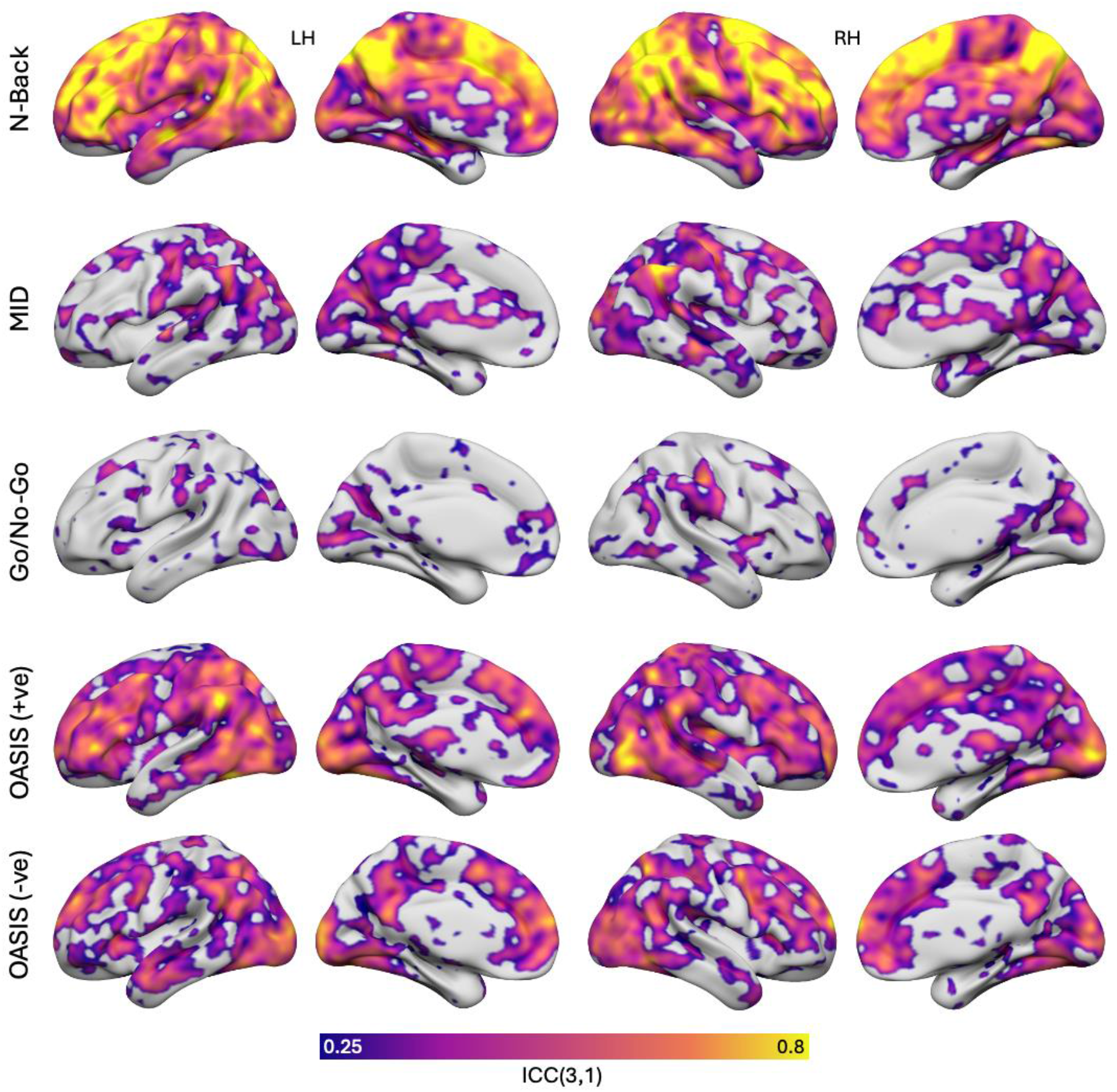
Whole brain voxelwise intraclass correlation coefficient (ICC(3,1)) maps (thresholded at 0.25) rendered on a smoothed cortical surface for the 6HMP pipeline and each task, where colour represents ICC values. OASIS (+ve) = OASIS positive > neutral and OASIS (-ve) = OASIS negative > neutral.

ICC values varied considerably across tasks but with comparatively smaller variation across preprocessing pipelines **(Figure 6)**. The N-back task demonstrated the highest reliability, with ICC values narrowly ranging from 0.423 (6HMP-aCompCor) to 0.490 (6HMP), suggesting an insensitivity to pipeline choice. These values fall within the “fair” range of ICC values (Cicchetti, 1994). The OASIS task showed ICC values ranging from 0.21 to 0.27, while both the MID and Go/No-Go tasks had uniformly low reliability (all ICCs < 0.15). See **Table S7** for full ICC results and associated confidence intervals.

**Figure 6.**
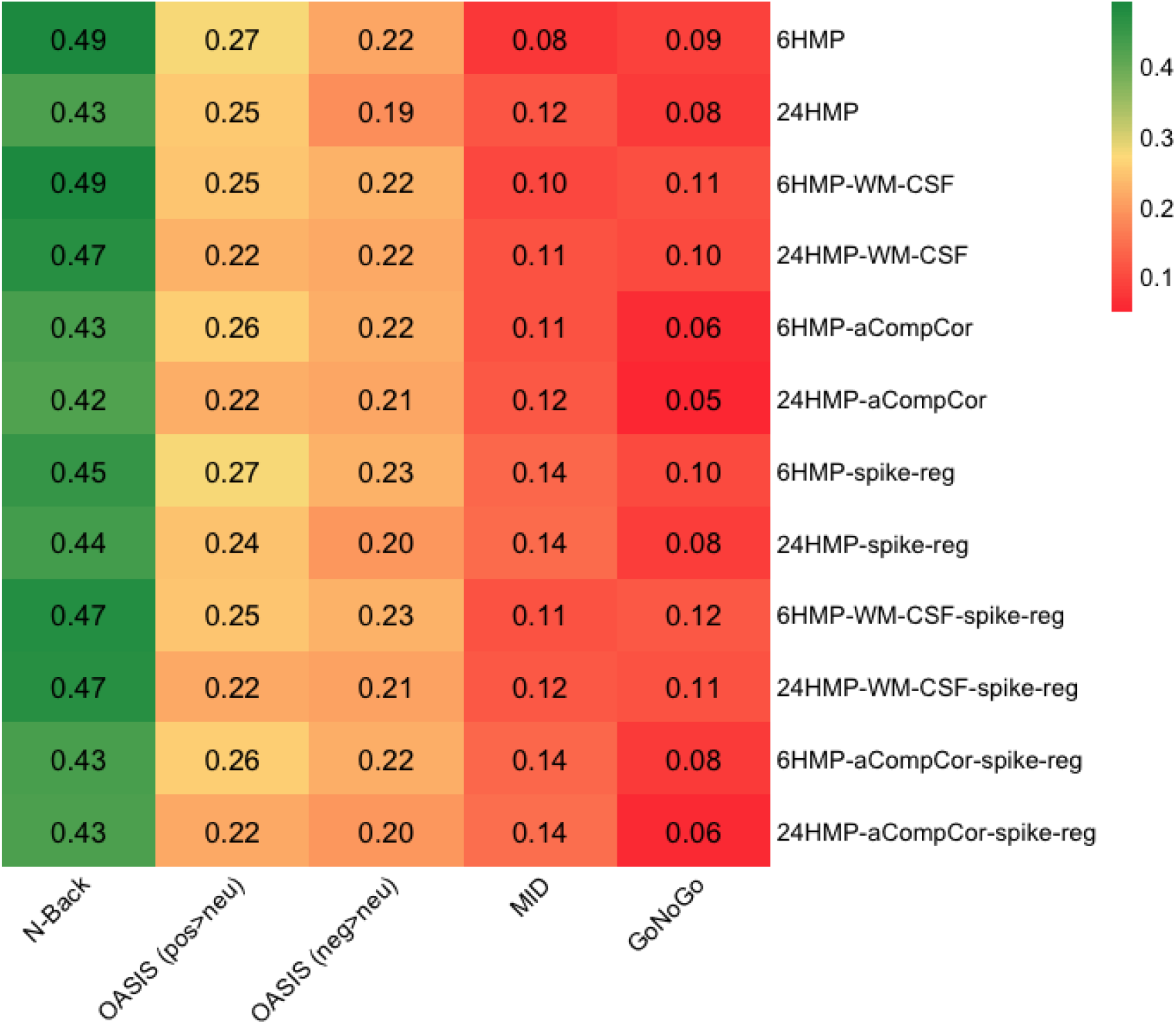
Heatmap of intraclass correlation coefficient (ICC) values from task-relevant ROI masks for each pipeline (rows) and task (columns). OASIS contrasts are positive>neutral and negative>neutral. Colour represents the mean ICC value, with better values in green and poorer values in red.

### Spearman’s correlation analyses

Spearman’s correlations were calculated to assess test-retest reliability of activation patterns within task-relevant regions between sessions (**Table 4**). Consistent with previous results, the N-back task showed excellent reliability with all values >0.95. The MID also showed excellent reliability with all values >0.8. The OASIS task values showed greater variability ranging between 0.54-0.76, with a similar result for the Go/No-Go task values (0.58-0.70).

**Table 4.** Spearman’s rank correlation coefficients between Session 1 and 2 group-level *Z*-statistic maps for each task and preprocessing pipeline.

|  | N-back | MID | Go/No-Go | OASIS (neg>neu) | OASIS (pos>neu) |
|---|---|---|---|---|---|
| **6HMP** | **0.966** | **0.893** | 0.588 | 0.694 | 0.586 |
| **24HMP** | 0.959 | 0.881 | 0.584 | 0.734 | 0.559 |
| **6HMP-WM-CSF** | 0.956 | 0.839 | 0.662 | 0.659 | **0.700** |
| **24HMP-WM-CSF** | 0.964 | 0.888 | 0.579 | 0.705 | 0.570 |
| **6HMP-aCompCor** | 0.953 | 0.820 | 0.666 | 0.666 | 0.696 |
| **24HMP-aCompCor** | 0.962 | 0.835 | 0.675 | 0.746 | 0.577 |
| **6HMP-spike-reg** | 0.960 | 0.831 | 0.680 | **0.759** | 0.562 |
| **24HMP-spike-reg** | 0.959 | 0.878 | 0.575 | 0.687 | 0.539 |
| **6HMP-WM-CSF-spike-reg** | 0.957 | 0.820 | 0.693 | 0.742 | 0.574 |
| **24HMP-WM-CSF-spike-reg** | 0.955 | 0.817 | **0.699** | 0.728 | 0.559 |
| **6HMP-aCompCor-spike-reg** | 0.954 | 0.814 | 0.652 | 0.638 | 0.697 |
| **24HMP-aCompCor-spike-reg** | 0.951 | 0.802 | 0.669 | 0.626 | 0.690 |

*Note*: Bold values indicate the highest coefficient per task.

## Discussion

This study examined how 12 different denoising strategies influenced the results and test-retest reliability of four common fMRI tasks using a constrained multiverse framework. All tasks produced a robust set of activation clusters consistent with previous studies using these tasks. These results serve to support the validity of task design, data acquisition, and the overall preprocessing and analysis approach. The measure of pipeline performance (FD-DVARS correlations) showed that the more complex pipelines removed more motion-related variance than the simpler pipelines (**Figure 1**), albeit at a greater cost in temporal degrees of freedom (**Figure S4, Table S6).** Despite this, denoising pipelines did not substantially alter the derived *Z-*statistic values (**Table 2, Figures S5-SG**). The spatial extent of clusters (number of suprathreshold voxels) was strongly influenced by pipeline in some tasks (notably the MID and the Go/No-Go tasks; **Figure 3**, **Table 2**). Reliability measures differed markedly across the four tasks, with the relative performance of denoising pipelines varying between tasks (**Figures 5-6**, **Tables 3-4**).

Spatial extents of suprathreshold clusters are a key measure in fMRI analyses (with more extensive clusters often regarded positively) and were substantially influenced by denoising pipeline selection (**Figure 3**, **Table 2**). In the N-back task, the pipeline choice had relatively little effect, which may reflect the high activation strength, large clusters, and robust task-evoked effects seen in this task. This may arise from the large network of cortical regions involved in working memory and/or the high cognitive load imposed by the task. Evidently, variations in the statistical modelling matter less when effects are strong. In the Go/No-Go task, pipelines incorporating tissue-based regressors (WM/CSF and aCompCor) were more favourable. The reverse was true for the MID where tissue-based regressors were associated with reduced voxel counts and pipelines only using head-motion parameters (6/24 head-motion regressors, spike regression) produced more extensive clusters. These differences may be related to distinct patterns of activation distribution in each task with greater sub-cortical involvement in the MID and more distributed cortical activation in the Go/No-Go task. In the OASIS task there were divergent pipeline effects at the level of individual task contrasts. Pipelines incorporating aCompCor regressors produced greater suprathreshold voxel counts in the positive > neutral contrast but produced fewer than other pipelines in the negative > neutral contrast. It is important to note that larger and more widespread clusters may not necessarily mean that results are higher quality. For example, in the MID task, results vary widely across pipelines, with extensive activation in some pipelines, beyond the regions typically associated with reward processing. This highlights a potential trade-off between sensitivity (obtaining large, robust clusters) and specificity (obtaining results specific to the particular process under investigation). In this case, the tissue-based regressors arguably provide less sensitive, but more highly specific results.

This highlights an issue with selection of analysis strategies for fMRI studies that include multiple tasks; there is unlikely to be a universally optimal pipeline that delivers robust performance across tasks (Jones et al., 2022). However, choosing a pipeline based on expected distribution of activation in the cortex and sub-cortex may be a useful strategy. Most troublingly, in the OASIS task there are divergent pipeline effects at the level of individual task contrasts, with pipelines including aCompCor regressors producing stronger results in the positive > neutral contrast, and other pipelines favouring the negative > neutral contrast. This suggests that pipeline selection can potentially bias outcomes even *within* a single task, depending on the contrasts examined.

Test-retest reliability remains a key challenge for fMRI studies and (based on the present results) more sophisticated denoising pipelines are unlikely to substantially improve reliability. This study adds to a growing body of literature (e.g. Demidenko et al., 2024; Elliott et al., 2020; Infantolino et al., 2018) in showing that fMRI measures can be reliable at the group level (Dice coefficients, Spearman’s correlations), while having limited reliability at the participant-level (ICC). While the overall results suggest that there is likely no universally optimal denoising method, they do provide a basis for making several recommendations. First, while adding spike/motion-censoring regressors to a model does reduce residual motion-related variance (assessed by the FD-DVARS measure), it seems to have relatively little effect on any of the sensitivity or reliability measures; the addition of spike regressors yields similar results to the equivalent pipeline without those regressors in all tasks. Given that it seems to confer little benefit and that it adds a variable (and potentially large) number of extra regressors to each first-level model, the current data would seem to discourage the use of the spike regression approach. Second, while exhaustive testing of different denoising methods for each dataset may be impractical, use of sensitivity analyses may be a useful approach. Researchers could select a primary denoising method but also repeat analyses with an additional pipeline to test sensitivity. Robust results across pipelines would provide greater confidence in observed effects. Alternatively, if the results vary, sensitivity itself may be important information. Third, reporting of denoising procedures should be both transparent and comprehensive. The choice of denoising can have a large impact on results; denoising should therefore be conceptualised as part of the analysis rather than as a relatively objective cleaning stage of preprocessing. Fourth, future efforts to investigate denoising approaches should ideally take a multidimensional approach, which is not dependent on any individual task, outcome measure, dataset, or study population. Benchmarking of denoising methods is complex, and our results are both task and outcome dependent. Previous work has also shown that single measures, such as the ICC, are not sufficient to characterise reliability (Gorgolewski et al., 2013) and has recommended the use of multiple measures when evaluating denoising pipelines (Zhai et al., 2026). Fifth, future investigations should seek to examine in what contexts particular methods perform well, rather than focusing on identifying an ‘optimal’ method. Investigations could examine how denoising approaches interact with head motion, task design, effect size, physiological factors, and subject populations, providing guidance for selecting the most appropriate analysis decisions for specific datasets. Sixth, more complex pipelines do not guarantee higher-quality results. For example, complex pipelines involving 24 head-motion parameters, aCompCor, and spike regression were often the least favourable across the different reliability metrics, while simple pipelines involving only six head motion parameters consistently performed better on these measures.

Across most measures used here, the most salient feature was the task itself, rather than the denoising pipeline applied. Consistent with previous work (Elliott et al., 2020), the N-back working memory task used here produced the most robust and reliable results. MID and Go/No-Go tasks performed poorly on reliability measures, and the OASIS task had intermediate performance. Previous work focused on replicability (distinct from reliability) also found that cognitive and emotional tasks were most replicable in small sample sizes, and a response inhibition task was the least replicable (Bossier et al., 2020). Also in line with previous work (e.g. Demidenko et al., 2024), we found that individual reliability measures (ICCs) were relatively poor across all tasks, whereas group-level measures (Dice coefficients, Spearman’s correlations) were substantially higher. The two block-design tasks used here (N-back, OASIS) outperformed event-related designs (Go/No-Go, MID) in reliability, consistent with literature showing that block-designs often show higher effect sizes than event-related designs (Chee et al., 2003; Tie et al., 2009). With sufficiently robust data (e.g. the N-back task), the exact processing and analysis pipeline used is unimportant. Researchers concerned about test-retest reliability should prioritise maximising experimental power in their studies by optimising task design (e.g. selecting block-designs where possible, and/or using longer tasks with more trials/blocks), using acquisition sequences with high signal-to-noise ratios (Wall et al., 2023), scanning each participant for longer on each task paradigm, and increasing participant numbers. All these *ex ante* considerations at the experimental design stage are likely to have a greater influence on reliability than any *post hoc* corrections applied at the preprocessing/analysis stage, no matter how sophisticated.

Similar arguments apply to the ‘multiverse’ analyses presented in **Figure 2** (see also visualisations of the largest/smallest activation maps in **Figure 4**). There is a recent trend toward conducting analyses with different denoising pipelines; for example, a recent mega-analysis paper (Girn et al., 2026) reported results using both aCompCor and ICA-AROMA (Pruim et al., 2015), as well as with and without global signal regression. Such examples may be motivated by either uncertainty about the optimal preprocessing strategy, or a desire to demonstrate the robustness of findings to methodological choices, particularly regarding contentious steps such as global signal regression (Murphy C Fox et al., 2017). In the present data, for tasks with little variability across the pipelines (e.g. N-back) the multiverse approach has quite limited utility, as the output of any individual pipeline closely approximates the consensus (averaged) result. For tasks with greater variability (e.g. Go/No-Go, MID), an across-pipeline consensus estimate may be more informative. However, these results suggest that a more appropriate solution would be to improve the experimental and task design to reduce this variability, rather than relying on multiverse analyses or any specific analysis pipeline. In practical terms, if all tasks were as robust as the N-back, preprocessing and denoising choices would have minimal impact, and multiverse analyses would also therefore be largely redundant.

This study has several limitations. First, some of the measures used (e.g. Dice coefficients, Spearman’s correlations) are most meaningfully defined at the group level, yielding a single summary value per pipeline. As a result, statistical comparison across pipelines on these measures is not possible. Second, these analyses were conducted on a single dataset of healthy individuals, limiting generalisability to other samples. Reliability estimates are strongly impacted by between-participant variance (which may be higher in clinical or developmental samples). Third, this “multiverse” analysis was restricted to variations in a single aspect of the analysis pipeline (confound regression; and only a subset of possible options). A wide range of alternative denoising approaches exist, including ICA-based methods such as ICA-AROMA (Pruim et al., 2015) and ICA-FIX (Salimi-Khorshidi et al., 2014), physiological noise correction approaches (Glover et al., 2000), and more recent novel methods (e.g. ‘rapidtide’; Aslan et al., 2025). Other aspects of the analysis pipeline (e.g. motion-correction method, details of temporal/spatial filtering) were held constant, although some of these features have been examined in previous work (Dafflon et al., 2022; Demidenko et al., 2024). Even this constrained approach required estimation of 2400 first-level models, made feasible by access to a high-performance computing system. An exhaustive evaluation of all possible fMRI pipeline variations would be impractical for most datasets, given the required computational and storage demands.

To conclude, this ‘multiverse’ analysis of denoising options across four commonly used task paradigms has demonstrated that test-retest reliability in fMRI is largely dependent on the task used (and the resulting effect sizes and robustness of the data). Block-design tasks are more robust and reliable than event-related tasks, and pipeline-related variability had more influence over derived effects when experimental power was lower. There may be no single optimal pipeline that achieves the best performance across different tasks. While constrained multiverse approaches may be useful, they are no substitute for well-powered, robust datasets, and well-designed tasks. Researchers who are concerned with test-retest reliability should prioritise maximising experimental power through optimisation of task design, acquisition parameters, and sample size.

## Data and Code Availability

The participants did not consent to open sharing of the primary data; however, it may be possible to share anonymised data to individual researchers/labs, on reasonable request, and following completion of a data-sharing agreement. Analysis code is available at https://github.com/saharuppal/task_based_multiverse_reliability_study.

## Author Contributions

**Sahar Uppal**: Methodology, Software, Formal analysis, Visualization, Writing - Original Draft, Writing - Review and Editing. **Elizabeth Davies**: Formal analysis, Visualization, Writing - Original Draft, Writing - Review and Editing**. Natalie Ertl**: Methodology, Investigation, Writing - Review and editing. **Lysia Demetriou**: Conceptualization, Methodology, Investigation, Writing - Review and editing. **Leon Fonville:** Writing - Review and Editing. **Matthew B Wall**: Conceptualization, Methodology, Software, Investigation, Visualization, Writing - Original Draft, Writing - Review and Editing, Supervision, Project administration.

## Acknowledgements

The authors would like to thank the clinical and operations staff at Perceptive Inc. (formerly ‘Invicro Ltd’) at the clinical imaging centre, Hammersmith Hospital for their help and support in conducting this project. We would also like to thank the staff at the Imperial College London High-Performance Computing research group for their support in the analysis phase.

