## Supplementary Material for "A test-retest reliability and multiverse analysis of denoising pipelines in four common functional MRI task paradigms"

*Full boiler-plate text produced by fMRIPrep:*

Results included in this manuscript come from preprocessing performed using *fMRIPrep* 24.1.1 (Esteban et al. (2019); Esteban et al. (2018); RRID:SCR_016216), which is based on *Nipype* 1.8.6 (K. Gorgolewski et al. (2011); K. J. Gorgolewski et al. (2018); RRID:SCR_002502).

### Anatomical data preprocessing

A total of 2 T1-weighted (T1w) images were found within the input BIDS dataset. Each T1w image was corrected for intensity non-uniformity (INU) with N4BiasFieldCorrection (Tustison et al. 2010), distributed with ANTs 2.5.3 (Avants et al. 2008, RRID:SCR_004757). The T1w-reference was then skull-stripped with a *Nipype* implementation of the antsBrainExtraction.sh workflow (from ANTs), using OASIS30ANTs as target template. Brain tissue segmentation of cerebrospinal fluid (CSF), white-matter (WM) and gray-matter (GM) was performed on the brain-extracted T1w using fast (FSL (version unknown), RRID:SCR_002823, Zhang, Brady, and Smith 2001). An anatomical T1w-reference map was computed after registration of 2 <module 'nipype.interfaces.image' from '/opt/conda/envs/fmriprep/lib/python3.11/site-packages/nipype/interfaces/image.py'> images (after INU-correction) using mri_robust_template (FreeSurfer 7.3.2, Reuter, Rosas, and Fischl 2010). Brain surfaces were reconstructed using recon-all (FreeSurfer 7.3.2, RRID:SCR_001847, Dale, Fischl, and Sereno 1999), and the brain mask estimated previously was refined with a custom variation of the method to reconcile ANTs-derived and FreeSurfer-derived segmentations of the cortical gray-matter of Mindboggle (RRID:SCR_002438, Klein et al. 2017). Volume-based spatial normalization to two standard spaces (MNI152NLin6Asym, MNI152NLin2009cAsym) was performed through nonlinear registration with antsRegistration (ANTs 2.5.3), using brain-extracted versions of both T1w reference and the T1w template. The following templates were were selected for spatial normalization and accessed with *TemplateFlow* (24.2.0, Ciric et al. 2022): *FSL's MNI ICBM 152 non-linear 6th Generation Asymmetric Average Brain Stereotaxic Registration Model* [Evans et al. (2012), RRID:SCR_002823; TemplateFlow ID: MNI152NLin6Asym], *ICBM 152 Nonlinear Asymmetrical template version 2009c* [Fonov et al. (2009), RRID:SCR_008796; TemplateFlow ID: MNI152NLin2009cAsym].

### Functional data preprocessing

For each of the 10 BOLD runs found per subject (across all tasks and sessions), the following preprocessing was performed. First, a reference volume was generated, using a custom methodology of *fMRIPrep*, for use in head motion correction. Head-motion parameters with respect to the BOLD reference (transformation matrices, and six corresponding rotation and translation parameters) are estimated before any spatiotemporal filtering using mcflirt (FSL , Jenkinson et al. 2002). The BOLD reference was then co-registered to the T1w reference using bbregister (FreeSurfer) which implements boundary-based registration (Greve and Fischl 2009). Co-registration was configured with six degrees of freedom. Several confounding time-series were calculated based on the *preprocessed BOLD*: framewise displacement (FD), DVARS and three region-wise global signals. FD was computed using two formulations following Power (absolute sum of relative motions, Power et al. (2014)) and Jenkinson (relative root mean square displacement between affines, Jenkinson et al. (2002)). FD and DVARS are calculated for each functional run, both using their implementations in *Nipype* (following the definitions by Power et al. 2014). The three global signals are extracted within the CSF, the WM, and the whole-brain masks. Additionally, a set of physiological regressors were extracted to allow for component-based noise correction (*CompCor*, Behzadi et al.

2007). Principal components are estimated after high-pass filtering the *preprocessed BOLD* time-series (using a discrete cosine filter with 128s cut-off) for the two *CompCor* variants: temporal (tCompCor) and anatomical (aCompCor). tCompCor components are then calculated from the top 2% variable voxels within the brain mask. For aCompCor, three probabilistic masks (CSF, WM and combined CSF+WM) are generated in anatomical space. The implementation differs from that of Behzadi et al. in that instead of eroding the masks by 2 pixels on BOLD space, a mask of pixels that likely contain a volume fraction of GM is subtracted from the aCompCor masks. This mask is obtained by dilating a GM mask extracted from the FreeSurfer's *aseg* segmentation, and it ensures components are not extracted from voxels containing a minimal fraction of GM. Finally, these masks are resampled into BOLD space and binarized by thresholding at 0.99 (as in the original implementation). Components are also calculated separately within the WM and CSF masks. For each CompCor decomposition, the *k* components with the largest singular values are retained, such that the retained components' time series are sufficient to explain 50 percent of variance across the nuisance mask (CSF, WM, combined, or temporal). The remaining components are dropped from consideration. The head-motion estimates calculated in the correction step were also placed within the corresponding confounds file. The confound time series derived from head motion estimates and global signals were expanded with the inclusion of temporal derivatives and quadratic terms for each (Satterthwaite et al. 2013). Frames that exceeded a threshold of 0.5 mm FD or 1.5 standardized DVARS were annotated as motion outliers. Additional nuisance timeseries are calculated by means of principal components analysis of the signal found within a thin band (*crown*) of voxels around the edge of the brain, as proposed by (Patriat, Reynolds, and Birn 2017). All resamplings can be performed with *a single interpolation step* by composing all the pertinent transformations (i.e. head-motion transform matrices, susceptibility distortion correction when available, and co-registrations to anatomical and output spaces). Gridded (volumetric) resamplings were performed using nitransforms, configured with cubic B-spline interpolation.

Many internal operations of *fMRIPrep* use *Nilearn* 0.10.4 (Abraham et al. 2014, RRID:SCR_001362), mostly within the functional processing workflow. For more details of the pipeline, see the section corresponding to workflows in *fMRIPrep*'s documentation.

**Copyright Waiver**

The above boilerplate text was automatically generated by fMRIPrep with the express intention that users should copy and paste this text into their manuscripts *unchanged*. It is released under the CC0 license.

**Figure S1.**

*Representative schematic of the four fMRI tasks used in this study. (A) Go/No-Go: participants responded to Go stimuli (G) and withheld responses for No-Go stimuli (N) (B) Monetary Inventive Delay (MID): participants responded to a target (circle) following a cue (square) indicating potentially monetary gain. (C) OASIS: participants viewed and rated a series of emotionally valenced images. (D) N-back: participants completed 0-back and 2-back working memory conditions, indicating whether each stimulus matched the target presented 0 or 2 items before.*

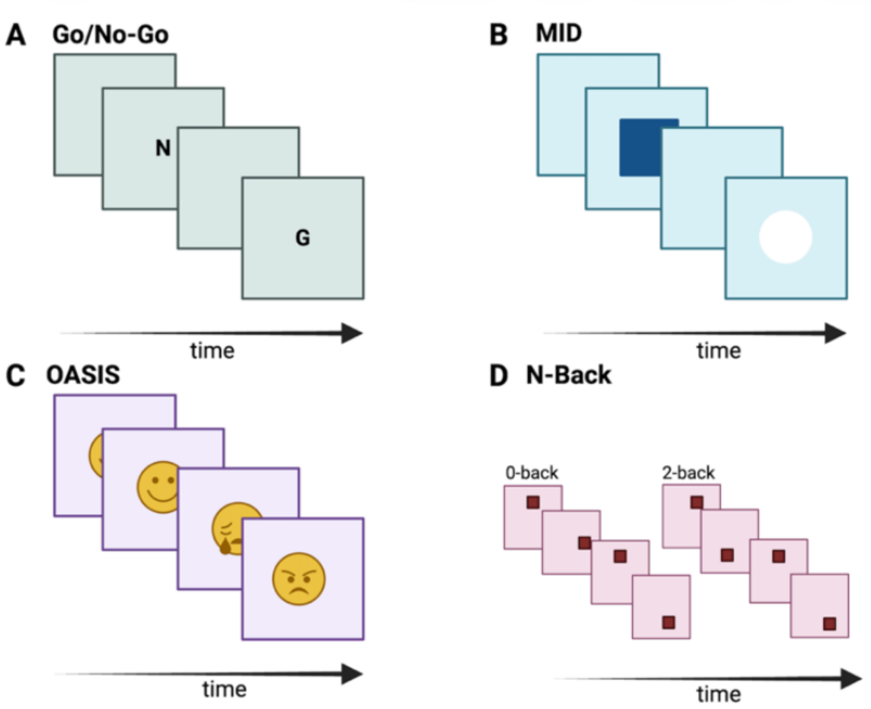


**Table S1.**

*Results of within-subject t-tests comparing head-motion (FD) across the two sessions, for each task.*

| Task | *t*-stat | *p*-value | $p_{FDR}$-value | Cohen’s *d* |
|---|---|---|---|---|
| N-back | -1.170 | 0.254 | 0.254 | 0.191 |
| MID | -0.089 | 0.930 | 0.951 | 0.019 |
| Go/No-Go | -2.760 | 0.011 | **0.043** | 0.429 |
| OASIS | 0.062 | 0.951 | 0.951 | -0.012 |

**Figure S2**

*Mean framewise displacement across all volumes for each task.*

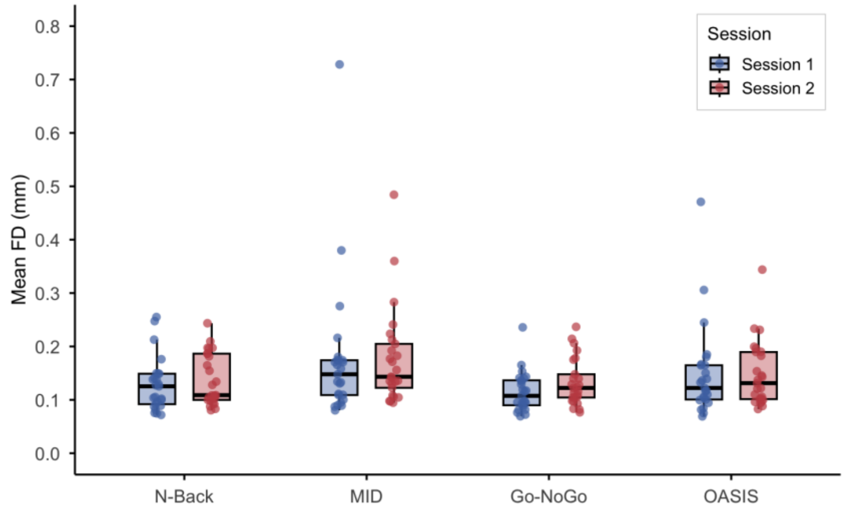


**Table S2**

*Number of motion-related spike regressors identified per subject, session and task.*

| Subject | Session | N-back | MID | Go/No-Go | OASIS |
|---|---|---|---|---|---|
| *Sub-001* | 1 | 52 | 38 | 13 | 80 |
| | 2 | 47 | 10 | 1 | 11 |
| *Sub-002* | 1 | 2 | 63 | 0 | 74 |
| | 2 | 166 | 73 | 77 | 71 |
| *Sub-003* | 1 | 0 | 6 | 0 | 20 |
| | 2 | 28 | 50 | 1 | 6 |
| *Sub-004* | 1 | 1 | 0 | 0 | 31 |
| | 2 | 0 | 32 | 0 | 0 |
| *Sub-005* | 1 | 0 | 4 | 0 | 1 |
| | 2 | 1 | 6 | 4 | 0 |
| *Sub-006* | 1 | 3 | 3 | 0 | 0 |
| | 2 | 0 | 18 | 2 | 2 |
| *Sub-007* | 1 | 21 | 2 | 7 | 7 |
| | 2 | 8 | 10 | 8 | 16 |
| *Sub-008* | 1 | 79 | 4 | 74 | 71 |
| | 2 | 70 | 20 | 22 | 0 |
| *Sub-009* | 1 | 66 | 93 | 129 | 19 |
| | 2 | 22 | 76 | 9 | 48 |
| *Sub-011* | 1 | 3 | 18 | 18 | 46 |
| | 2 | 19 | 45 | 42 | 2 |
| *Sub-012* | 1 | 33 | 136 | 75 | 0 |
| | 2 | 35 | 60 | 76 | 17 |
| *Sub-013* | 1 | 4 | 59 | 39 | 73 |
| | 2 | 65 | 48 | 18 | 84 |
| *Sub-014* | 1 | 129 | 44 | 252 | 75 |
| | 2 | 59 | 47 | 79 | 59 |

| | | | | | |
|---|---|---|---|---|---|
| *Sub-015* | 1 | 22 | 73 | 14 | 29 |
| | 2 | 16 | 16 | 28 | 11 |
| *Sub-016* | 1 | 7 | 13 | 42 | 6 |
| | 2 | 25 | 4 | 28 | 13 |
| *Sub-017* | 1 | 30 | 198 | 25 | 48 |
| | 2 | 100 | 125 | 54 | 41 |
| *Sub-018* | 1 | 0 | 0 | 0 | 0 |
| | 2 | 0 | 25 | 12 | 0 |
| *Sub-019* | 1 | 18 | 21 | 7 | 15 |
| | 2 | 22 | 66 | 67 | 37 |
| *Sub-020* | 1 | 5 | 0 | 1 | 2 |
| | 2 | 0 | 1 | 13 | 0 |
| *Sub-021* | 1 | 98 | 103 | 26 | 39 |
| | 2 | 51 | 109 | 16 | 31 |
| *Sub-022* | 1 | 3 | 0 | 32 | 0 |
| | 2 | 0 | 74 | 77 | 27 |
| *Sub-024* | 1 | 55 | 65 | 30 | 26 |
| | 2 | 72 | 147 | 14 | 39 |
| *Sub-025* | 1 | 35 | 26 | 24 | 44 |
| | 2 | 28 | 78 | 33 | 34 |
| *Sub-026* | 1 | 46 | 79 | 41 | 50 |
| | 2 | 38 | 77 | 33 | 32 |
| *Sub-027* | 1 | 4 | 27 | 22 | 1 |
| | 2 | 33 | 52 | 3 | 6 |

*Note:* a single motion-related spike regressor represents a single volume in which the framewise displacement of the participant exceeded 0.5 mm.

**Figure S3**

*Spearman correlations between mean FD and mean Z-statistic across pipelines and task.*

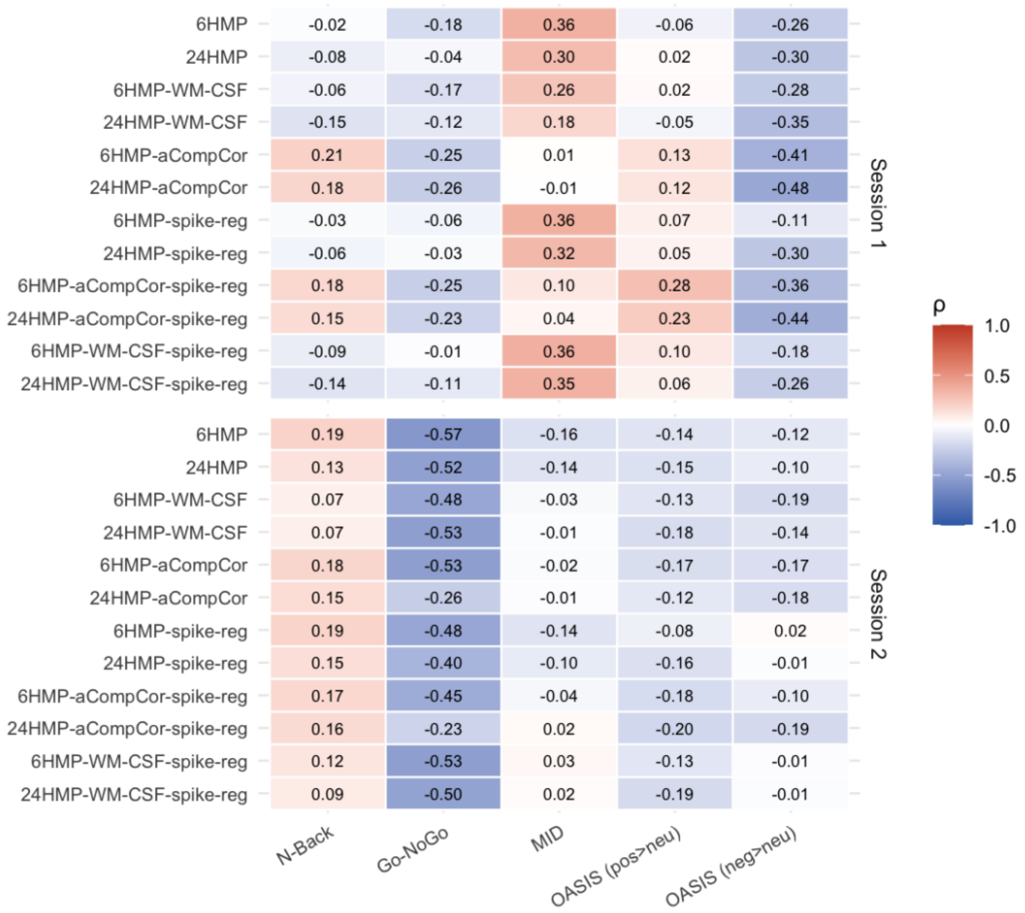


*Note*: $p_{(FDR)}<0.05$. Top plot = Session 1, bottom plot = Session 2. There were no significant correlations at the corrected threshold.

**Table S3.**

*Summary statistics for Pearson correlation between task explanatory variables (EV) and framewise displacement.*

| Task | EV | Median r | IQR | | Fisher Mean z | Fisher Mean r |
|---|---|---|---|---|---|---|
| Go/No-Go | Go | -0.045 | [-0.210, | 0.091] | -0.047 | -0.047 |
| Go/No-Go | No-Go | -0.001 | [-0.134, | 0.135] | 0.001 | 0.001 |
| MID | Neutral | 0.025 | [-0.129, | 0.118] | 0.016 | 0.016 |
| MID | Win | 0.065 | [-0.113, | 0.234] | 0.071 | 0.071 |
| N-back | 0-back | 0.000 | [-0.178, | 0.273] | 0.000 | 0.000 |
| N-back | 2-back | -0.081 | [-0.275, | 0.092] | -0.093 | -0.093 |
| OASIS | Positive | -0.018 | [-0.236, | 0.190] | -0.015 | -0.015 |
| OASIS | Negative | -0.010 | [-0.217, | 0.193] | -0.012 | -0.012 |
| OASIS | Neutral | 0.020 | [-0.156, | 0.184] | 0.017 | 0.017 |

**Table S4**

*Median spearman rho of framewise displacement and DVARS of the unexplained residuals after nuisance regression calculated for each pipeline within task.*

| Pipeline | N-back | MID | Go/No-Go | OASIS |
|---|---|---|---|---|
| 6HMP | 0.133 | 0.336 | 0.106 | 0.180 |
| 24HMP | 0.044 | 0.273 | **-0.003** | 0.066 |
| 6HMP-WM-CSF | 0.138 | 0.341 | 0.125 | 0.176 |
| 24HMP-WM-CSF | 0.042 | 0.271 | 0.005 | 0.076 |
| 6HMP-aCompCor | 0.220 | 0.367 | 0.184 | 0.227 |
| 24HMP-aCompCor | 0.126 | 0.334 | 0.085 | 0.122 |
| 6HMP-spike-reg | 0.080 | 0.266 | 0.075 | 0.069 |
| 24HMP-spike-reg | **-0.017** | **0.152** | -0.040 | -0.017 |
| 6HMP-WM-CSF-spike-reg | 0.071 | 0.265 | 0.069 | 0.087 |
| 24HMP-WM-CSF-spike-reg | -0.044 | 0.159 | -0.006 | **0.006** |
| 6HMP-aCompCor-spike-reg | 0.137 | 0.319 | 0.172 | 0.173 |
| 24HMP-aCompCor-spike-reg | 0.069 | 0.249 | 0.029 | 0.051 |

*Note*: Rho values for pipelines producing the smallest association between these variables are in bold and columns are ordered in order of smallest to largest magnitudes.

**Table S5**

*Results of overall Spearman's rho FD-DVARS of pipelines ranked across task, sessions and subjects.*

| Pipeline | Median rank | % runs ranked 1st | % runs top 3 | % runs bottom 3 |
|---|---|---|---|---|
| 6HMP | 10 | 1 | 5 | 52 |
| 24HMP | 4 | 5 | 28 | 3 |
| 6HMP-WM-CSF | 10 | 0 | 2 | 55 |
| 24HMP-WM-CSF | 5 | 0 | 20 | 1 |
| 6HMP-aCompCor | 12 | 0 | 0 | 97 |
| 24HMP-aCompCor | 8 | 0 | 9 | 25 |
| 6HMP-spike-reg | 6 | 5 | 19 | 2 |
| 24HMP-spike-reg | 2 | 38 | 84 | 0 |
| 6HMP-WM-CSF-spike-reg | 6 | 2 | 12 | 5 |
| 24HMP-WM-CSF-spike-reg | 2 | 29 | 78 | 0 |
| 6HMP-aCompCor-spike-reg | 10 | 0 | 0 | 53 |
| 24HMP-aCompCor-spike-reg | 5 | 17 | 40 | 4 |

**Figure S4**

*Temporal degrees of freedom (tDOF) loss across the 12 pipelines, averaged across subjects and all four tasks.*

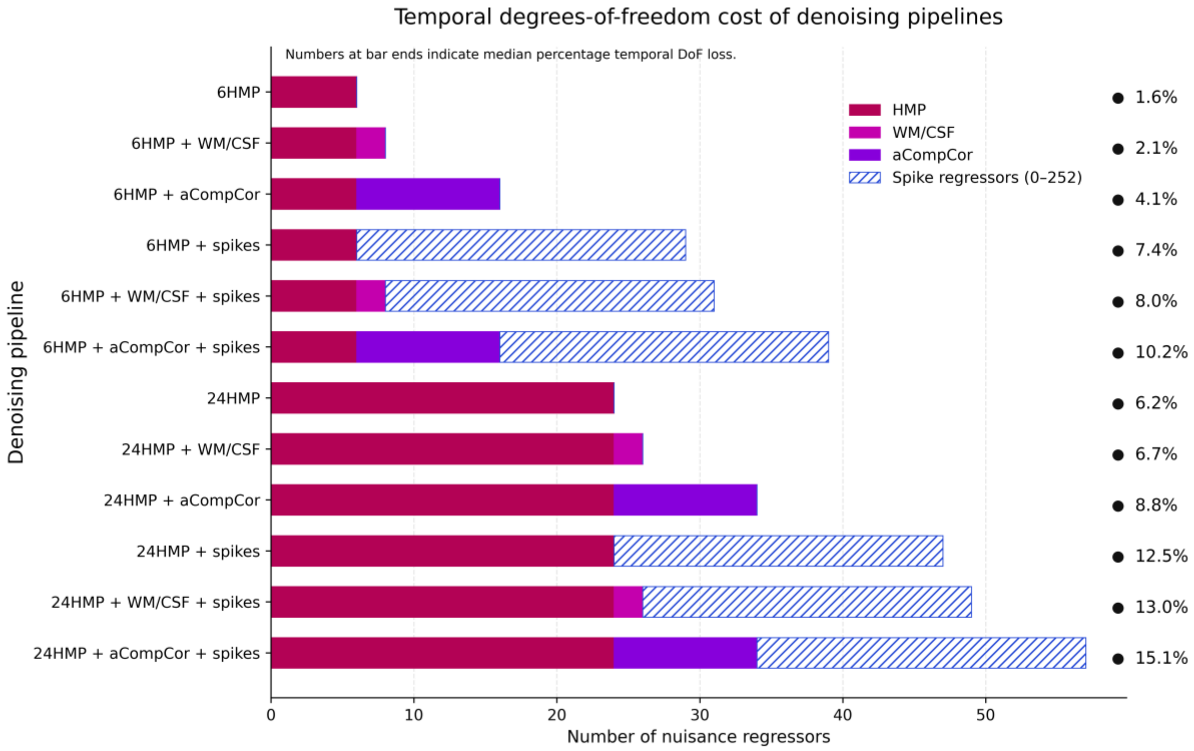


**Table S6**

*Results of temporal degrees of freedom loss pooled across subjects and task per pipeline.*

| Pipeline | **Pooled Median (%)** | **[IQR]** | **Min, Max (%)** |
|---|---|---|---|
| 6HMP | **1.55 %** | **[0.24]** | 1.23, 1.95 |
| 24HMP | 6.22 % | [0.94] | 4.92, 7.79 |
| 6HMP-WM-CSF | 2.07 % | [0.32] | 1.64, 2.60 |
| 24HMP-WM-CSF | 6.74 % | [1.02] | 5.33, 8.44 |
| 6HMP-aCompCor | 4.14 % | [0.63] | 3.28, 5.19 |
| 24HMP-aCompCor | 8.81 % | [1.34] | 6.97, 11.04 |
| 6HMP-spike-reg | 7.38 % | [12.32] | 1.23, 66.49 |
| 24HMP-spike-reg | 12.50 % | [12.08] | 4.92, 71.13 |
| 6HMP-WM-CSF-spike-reg | 7.96 % | [12.31] | 1.64, 67.01 |
| 24HMP-WM-CSF-spike-reg | 13.00 % | [11.91] | 5.33, 71.65 |
| 6HMP-aCompCor-spike-reg | 10.19 % | [12.56] | 3.28, 69.07 |
| 24HMP-aCompCor-spike-reg | 15.08 % | [11.53] | 6.97, 73.71 |

**Figure S5.**

*Mosaic of standard template slices (MNI152) overlain with significant clusters from the N-back task across the 12 pipelines. Thresholded at Z = 3.1,* $p_{(cluster\ corrected)} < 0.05$.

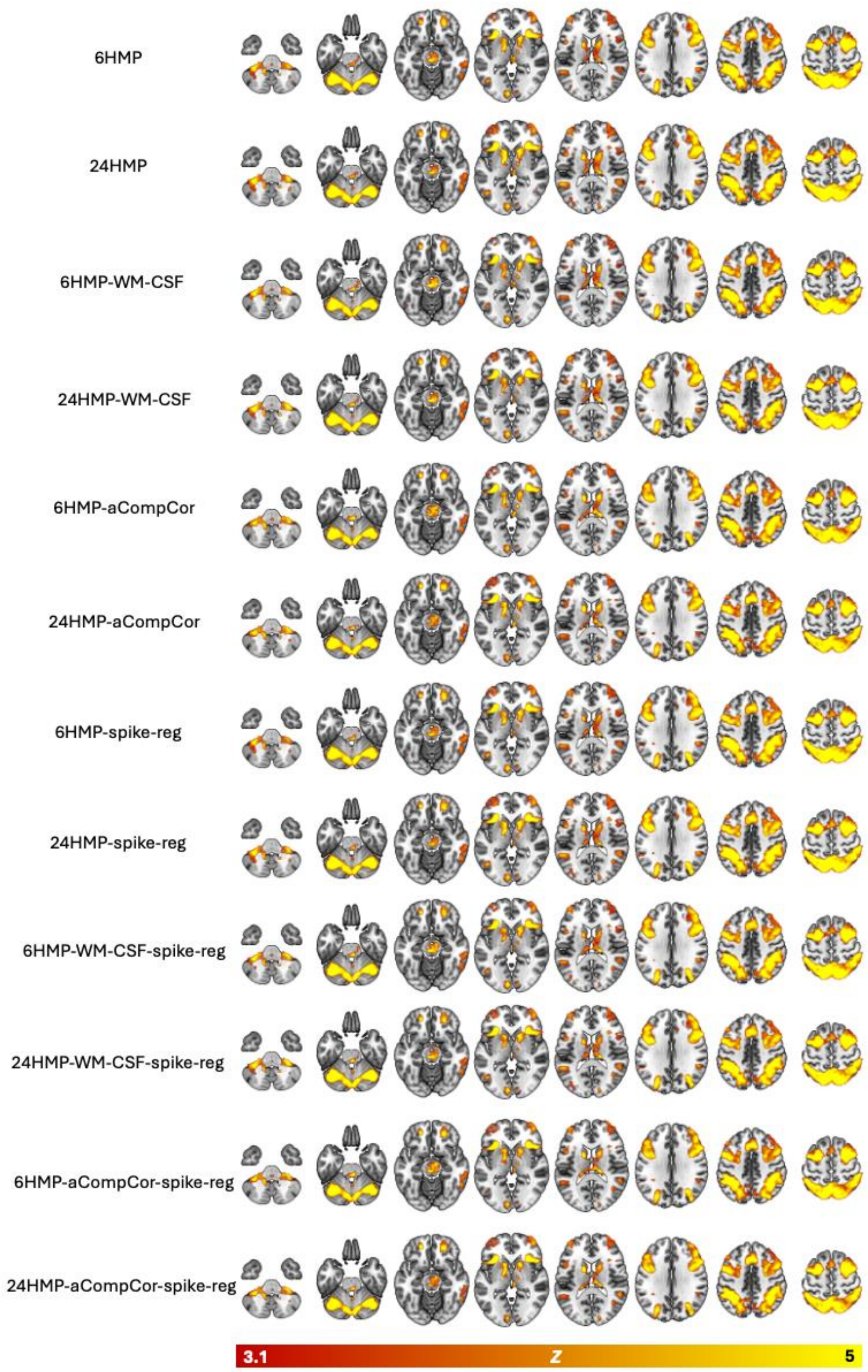

**Figure S6.**

*Mosaic of standard template slices (MNI152) overlain with significant clusters from the MID task across the 12 pipelines. Thresholded at* $Z = 3.1$, $p_{(cluster\ corrected)} < 0.05$.

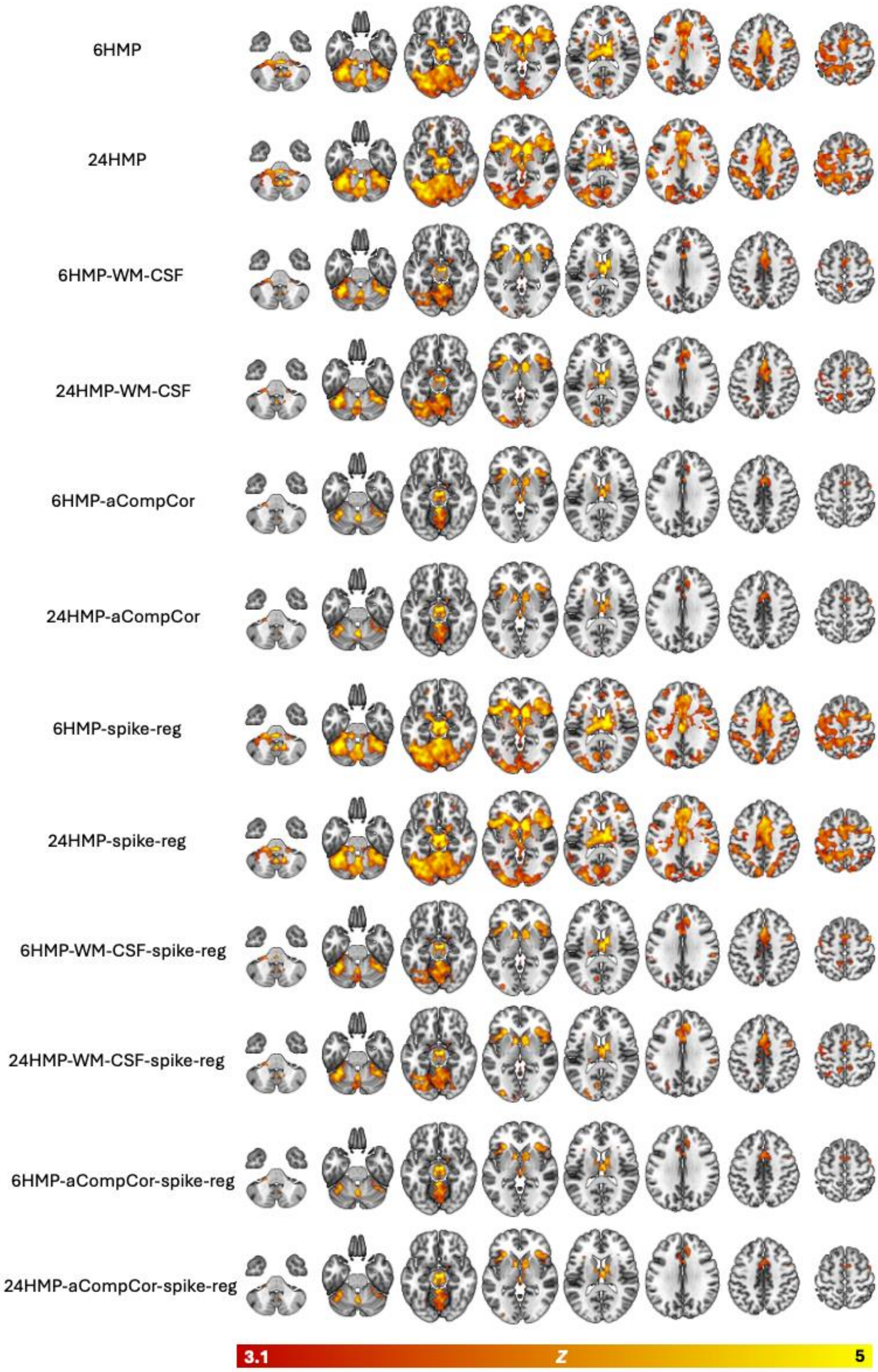

**Figure S7**

*Mosaic of standard template slices (MNI152) overlain with significant clusters from the Go/No-Go task across the 12 pipelines. Thresholded at $Z = 3.1$, $p_{(cluster\ corrected)} < 0.05$.*

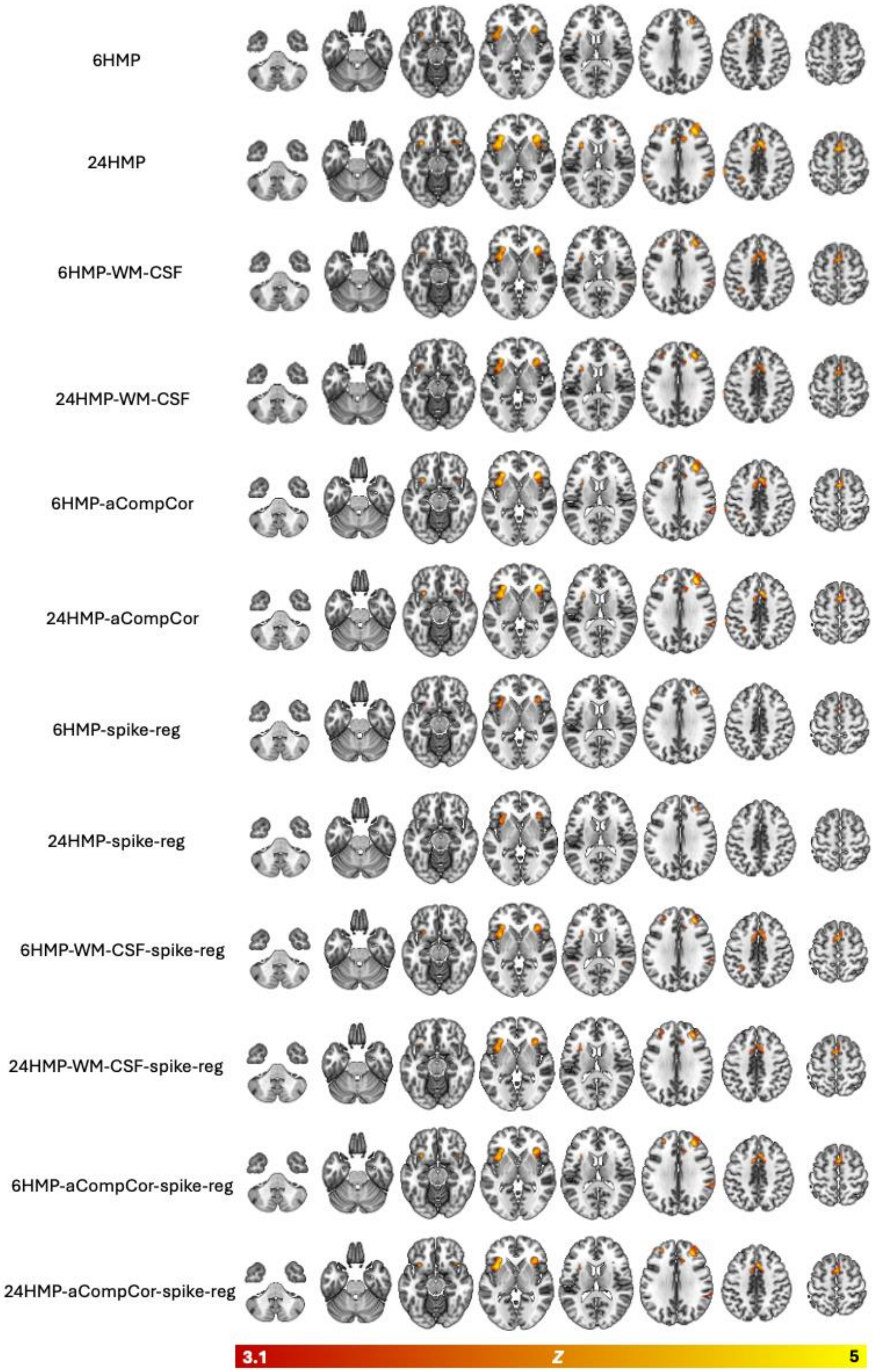

**Figure S8**

*Mosaic of standard template slices (MNI152) overlain with significant clusters from the OASIS (positive > neutral) task across the 12 pipelines. Thresholded at Z = 3.1, $p_{(cluster\ corrected)} < 0.05$.*

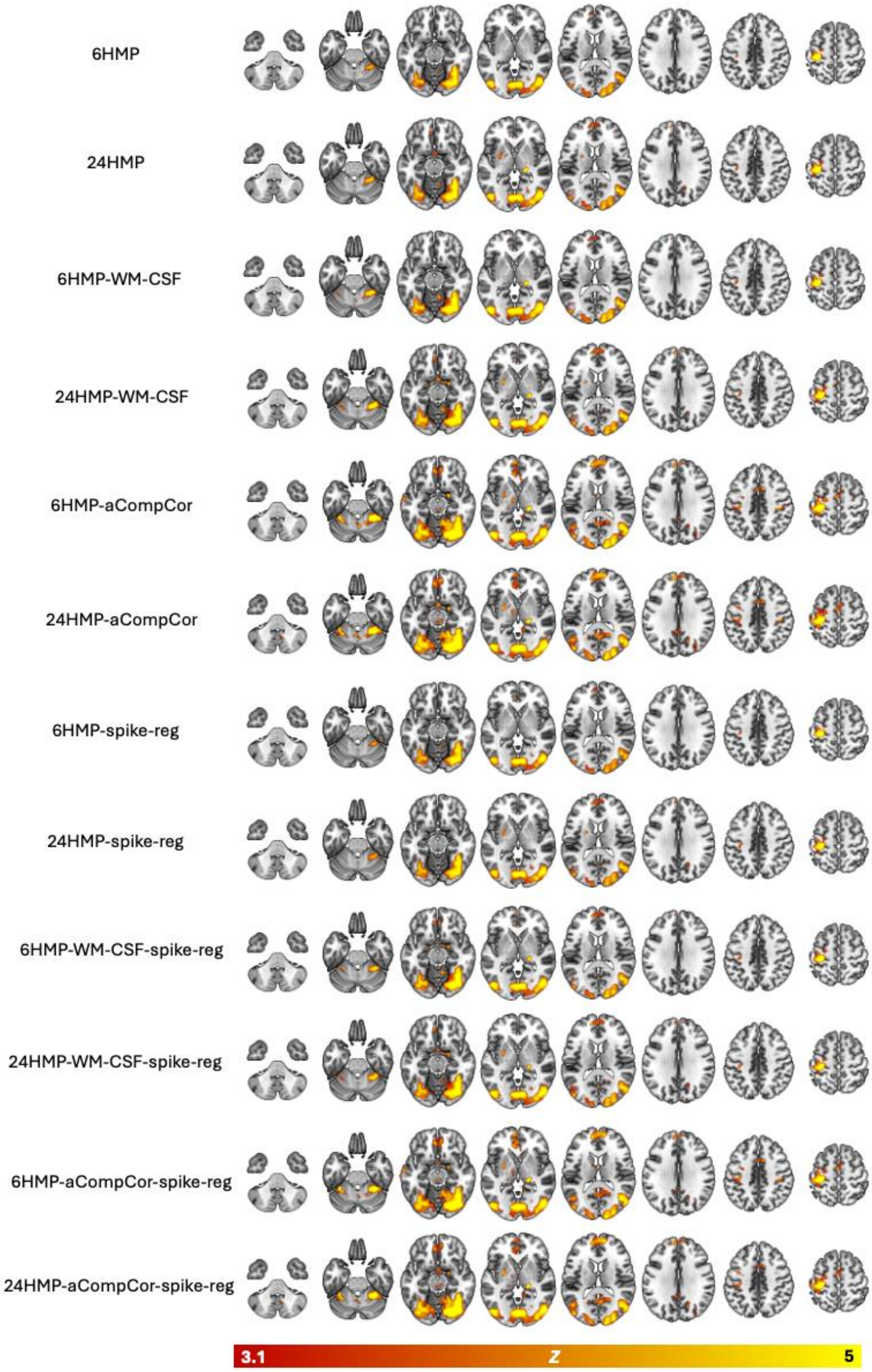

**Figure S9**

*Mosaic of standard template slices (MNI152) overlain with significant clusters from the OASIS (negative > neutral) task across the 12 pipelines. Thresholded at Z = 3.1, $p_{(cluster\ corrected)} < 0.05$.*

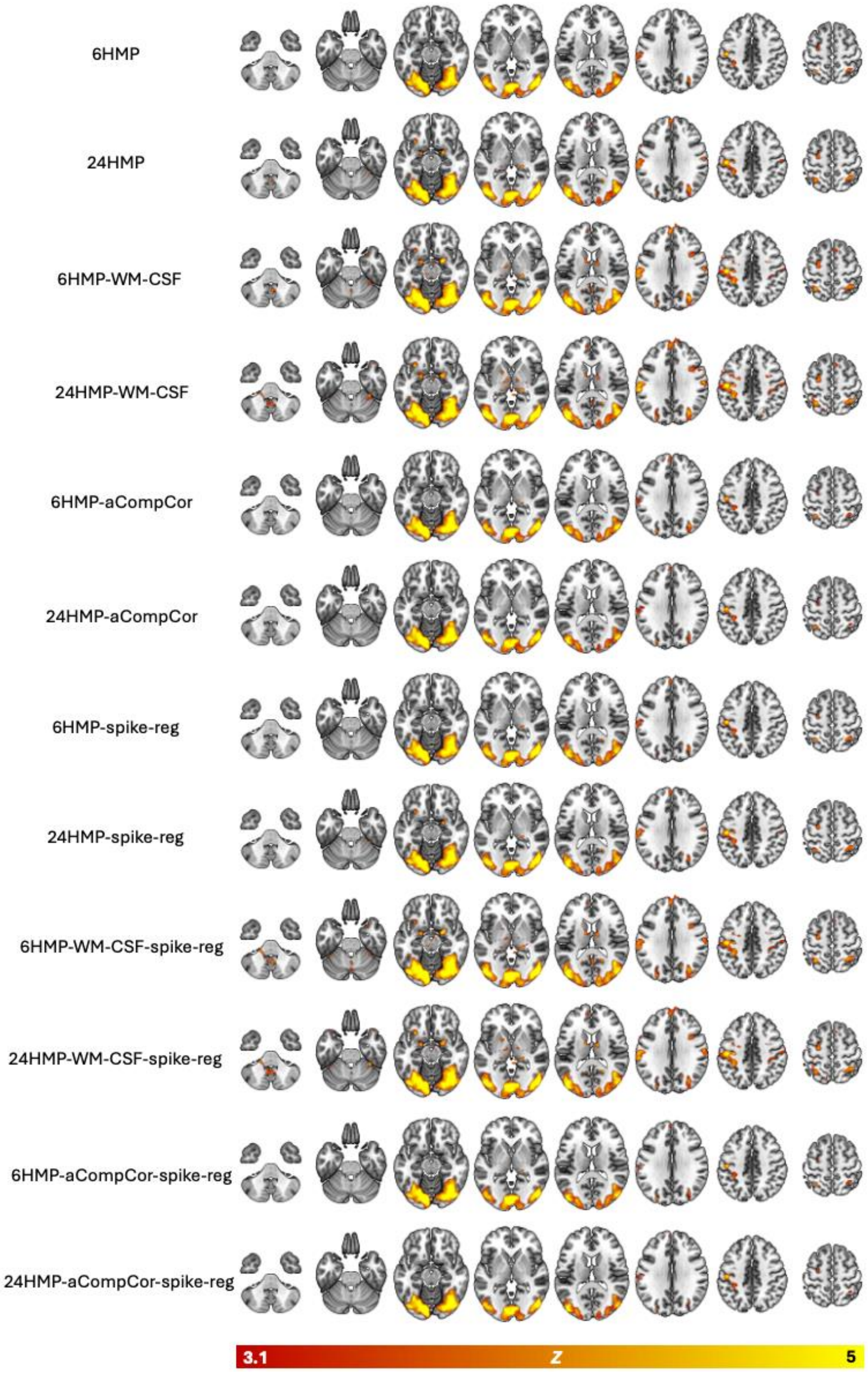

**Figure S10**

*Masks used for regional ICC extraction, displayed as axial slices overlaid on the MNI152 standard brain. These were generated using meta-analytic data and downloaded from [https://neurosynth.org/](https://neurosynth.org/) using relevant terms for each task: (A) 'Inhibitory control' (Go/No-Go) (B) 'Monetary' (MID task) (C) 'Working memory' (N-Back task) and (D) 'Emotional' (OASIS).*

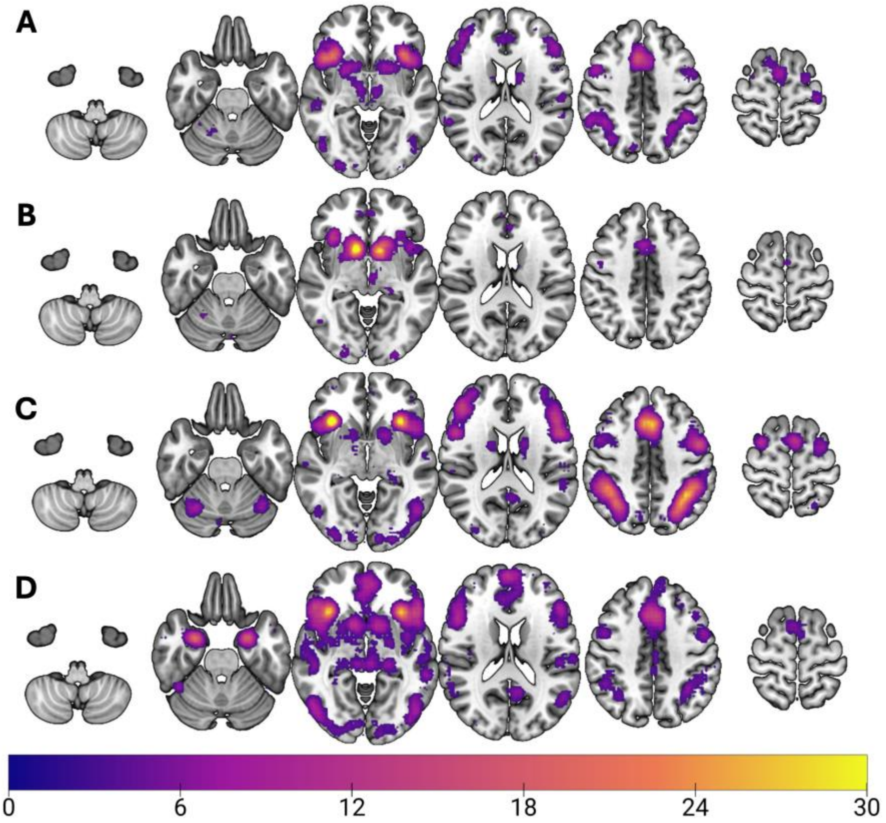

**Table S7**

*ICC (3,1) estimates and 95% confidence intervals for each task and preprocessing pipeline, derived using the masks in the previous figure.*

| | N-back | MID | Go/No-Go | OASIS (pos>neu) | OASIS (neg>neu) |
|---|---|---|---|---|---|
| 6HMP | **0.490 (0.486, 0.493)** | 0.077 (0.073, 0.081) | 0.087 (0.085, 0.090) | 0.273 (0.271, 0.276) | 0.215 (0.213, 0.218) |
| 24 HMP | 0.427 (0.424, 0.430) | 0.116 (0.111, 0.120) | 0.079 (0.077, 0.082) | 0.252 (0.249, 0.254) | 0.186 (0.184, 0.188) |
| 6HMP-WM-CSF | 0.488 (0.484, 0.491) | 0.098 (0.094, 0.103) | 0.107 (0.105, 0.110) | 0.248 (0.245, 0.250) | 0.225 (0.222, 0.227) |
| 24HMP-WM-CSF | 0.472 (0.468, 0.475) | 0.109 (0.105, 0.114) | 0.104 (0.101, 0.106) | 0.225 (0.222. 0.227) | 0.216 (0.214, 0.218) |
| 6HMP-aCompCor | 0.431 (0.427, 0.435) | 0.112 (0.108, 0.118) | 0.065 (0.063, 0.067) | 0.258 (0.256, 0.261) | 0.222 (0.219, 0.224) |
| 24HMP-aCompCor | 0.423 (0.419, 0.426) | 0.121 (0.116, 0.126) | 0.052 (0.049, 0.054) | 0.222 (0.219, 0.224) | 0.211 (0.209,214) |
| 6HMP-spike-reg | 0.454 (0.450-0.457) | 0.139 (0.134, 0.144) | 0.102 (0.100, 0.105) | **0.274 (0.272, 0.277)** | 0.226 (0.224, 0.229) |
| 24HMP-spike-reg | 0.436 (0.433, 0.440) | 0.143 (0.139, 0.148) | 0.085 (0.082, 0.087) | 0.245 (0.243, 0.247) | 0.195 (0.193, 0.198) |
| 6HMP-WM-CSF-spike-reg | 0.474 (0.470, 0.477 | 0.110 (0.106, 0.115) | **0.121 (0.118, 0.123)** | 0.247 (0.244, 0.249) | **0.227 (0.225- 0.230**) |
| 24HMP-WM-CSF-spike-reg | 0.486 (0.464, 0.471) | 0.119 (0.115, 0.124) | 0.113 (0.111, 0.116) | 0.217 (0.215, 0.220) | 0.208 (0.205, 0.210) |
| 6HMP-aCompCor-spike-reg | 0.429 (0.425, 0.432) | 0.140 (0.135, 0.145) | 0.081 (0.078, 0.083) | 0.259 (0.256, 0.262) | 0.223 (0.220-0.225) |
| 24HMP-aCompCor-spike-reg | 0.425 (0.422, 0.429) | **0.145 (0.140, 0.150)** | 0.060 (0.058, 0.062) | 0.220 (0.217, 0.222) | 0.196 (0.194, 0.198) |

*Note*: Bold values indicate the highest performing pipeline per task.
